# A retrieved context model of the emotional modulation of memory

**DOI:** 10.1101/175653

**Authors:** Deborah Talmi, Lynn J. Lohnas, Nathaniel D. Daw

**Author notes:** **Address for correspondence** Deborah Talmi, Division of neuroscience and experimental psychology, School of Biological Sciences, University of Manchester, Manchester, UK, M139PL.

## Abstract

Emotion enhances episodic memory, an effect thought to be an adaptation to prioritise the memories that best serve evolutionary fitness. But viewing this effect largely in terms of prioritising what to encode or consolidate neglects broader rational considerations about what sorts of associations should be formed at encoding, and which should be retrieved later. Although neurobiological investigations have provided many mechanistic clues about how emotional arousal modulates item memory, these effects have not been wholly integrated with the cognitive and computational neuroscience of memory more generally.

Here we apply the Context Maintenance and Retrieval Model (CMR, Polyn, Norman & Kahana, 2009) to this problem by extending it to describe the way people may represent and process emotional information. A number of ways to operationalise the effect of emotion were tested. The winning emotional CMR (eCMR) model reconceptualises emotional memory effects as arising from the modulation of a process by which memories become bound to ever-changing temporal and emotional contexts. eCMR provides a good qualitative fit for the emotional list-composition effect and the emotional oddball effect, illuminating how these effects are jointly determined by the interplay of encoding and retrieval processes. eCMR explains the increased advantage of emotional memories in delayed memory tests through the limited ability of retrieval to reinstate the temporal context of encoding.

By leveraging the rich tradition of temporal context models, eCMR helps integrate existing effects of emotion and provides a powerful tool to test mechanisms by which emotion affects memory in a broad range of paradigms.

## 1. Introduction

There is tremendous interest in the effect of emotional arousal on episodic memory. The literature generally agrees that moderate emotional arousal enhances item memory (Cahill & McGaugh, 1998; LaBar & Cabeza, 2006; Yonelinas & Ritchey, 2015). However, the circumstances in which it does so, and the mechanisms and models by which these effects are understood, might look unusual to a student of memory more generally. The mainstream human memory literature has traditionally tested memory within minutes of encoding because most functional manipulations do not dissociate immediate and delayed memory performance. By contrast, the research on emotional memory focuses on factors that influence delayed memory, and those are interpreted mostly in terms of prioritised storage. One reason for the appeal of that focus is that it suggests an interpretation in terms of the broader purpose of memory in guiding behaviour, and connections to an emerging set of rational, decision-theoretic accounts of the allocation of limited cognitive resources. In particular, emotionally-charged memories may be most relevant to fitness-relevant decisions later (Boureau, Sokol-Hessner, & Daw, 2015; Gershman & Daw, 2017), and therefore their maintenance should be prioritised in the face of limited memory capacity. However, we argue this view is incomplete from both an empirical and a normative perspective.

Empirically, current models of emotional memory fail to predict when enhancement would be exhibited and when it would not. In particular, simple prioritization fails to account for situations in which emotion does *not* benefit memory, such as in free recall tests for pure lists of emotional and neutral stimuli (the emotional list-composition effect, described below); in immediate recognition tests for mixed lists of emotional and neutral stimuli (Dougal & Rotello, 2007; Sharot & Yonelinas, 2008); and in tests of associative memory (Bisby, Horner, Hørlyck, & Burgess, 2016; Madan, Caplan, Lau, & Fujiwara, 2012; Madan, Fujiwara, Caplan, & Sommer, 2017). Furthermore, while the emotional memory literature has so far marginalised the influence of the retrieval test, in other areas of memory there has been a recent emphasis on retrieved context models of memory, which emphasise a role for associations between items and context during both encoding and retrieval (Howard & Kahana, 2002a; Lohnas, Polyn, & Kahana, 2015; Polyn, Norman, & Kahana, 2009; Sederberg, Howard, & Kahana, 2008). ‘Brain states’ that change gradually with time (Manns, Howard, & Eichenbaum, 2007) are thought to provide the neurobiological substrate that implements the temporal context in such models, a view supported by recent evidence that temporal overlap functionally links neural representations of separate events (Cai et al., 2016; Rashid et al., 2016).

Apart from addressing these empirical considerations, retrieved context models suggest a more nuanced normative interpretation. Instead of simply ranking which individual items should be prioritized in isolation, the retrieved context models speak to what sorts of memory structures should be built by encoding associations among these items. Indeed, these associations themselves have a clear normative interpretation in that the item-context associations at the heart of retrieved context models mathematically correspond to a particular sort of world model for guiding utility-maximizing future choices (Gershman, Moore, Todd, Norman, & Sederberg, 2012). Retrieved context models also shed light on the distinct question of how memories will be prioritized for *retrieval* given the goals of the test context (DuBrow, Rouhani, Niv, & Norman, 2017). In decision making, such modulation of retrieval will promote consideration of particular (emotionally charged) outcomes of candidate actions, and are reminiscent of mechanisms independently proposed in the decision-making literature for rationally prioritized evaluation (Cushman & Morris, 2015; Huys et al., 2012; Lieder, Griffiths, & Hsu, 2018).

Here we reinterpret emotional memory effects in the framework of retrieved context models with the aim to explain how emotion enhances episodic memory through their already-established mechanisms, thus bringing the emotional memory literature more closely into the fold of the mainstream memory literature. Our aims are, first, to consider systematically the different ways in which emotion can be operationalised within the Context Maintenance and Retrieval Model (CMR); and second, to identify which constellation of mechanisms allow the extended model, emotional CMR (eCMR), to capture qualitatively key effects of episodic memory for emotionally arousing items.

### 1.1 Existing models of emotional memory

The focus of the literature on emotional memory on delayed effects of emotion stems partly from the social value of understanding memory for key life events - experiences that define us as people (graduating from school, winning a competition) and as community members (weddings, funerals, flash-bulb memories for culturally important events). This remained a focus in the laboratory, where emotional experiences were operationalised by presenting participants with emotional words, pictures (FIGURE 1), stories and film clips, and testing memory for them 24 hours to two weeks later. Another contributing factor for the focus on delayed memory is that the dominant model for emotional memory, the modulated-consolidation model (Cahill & McGaugh, 1998; McGaugh et al., 2000), is concerned with effects that manifest themselves after a few hours, but not before. Both the Modulated-consolidation model and the Emotional Binding account (Yonelinas & Ritchey, 2015) attribute all of the explanatory power to processes that occur in the hours and days after stimulus encoding is completed. The former model suggests that emotional arousal directly modulates the consolidation of memory traces, a process that can be triggered when emotional stimuli are encoded but equally when arousal is experienced soon afterwards. The latter account suggests that the arousal experienced at encoding results in attenuated forgetting because of reduced interference from less-common emotional experiences, without influencing consolidation. GANE (Mather, Clewett, Sakaki, & Harley, 2015) also proposes a neurobiological mechanism for the effect of arousal and arousal-attention interactions on protein synthesis after encoding is completed. All of these models fare well in accounting for the oft-cited finding that the effect of emotion on memory is more robust in delayed, compared to immediate, memory tests (reviewed in Yonelinas & Ritchey, 2015 and earlier by Craik & Blankenstein, 1975) and for the effects of stressors and neuromodulators on memory for pre-stress experiences (Shields, Sazma, McCullough, & Yonelinas, 2017).

**FIGURE 1.**
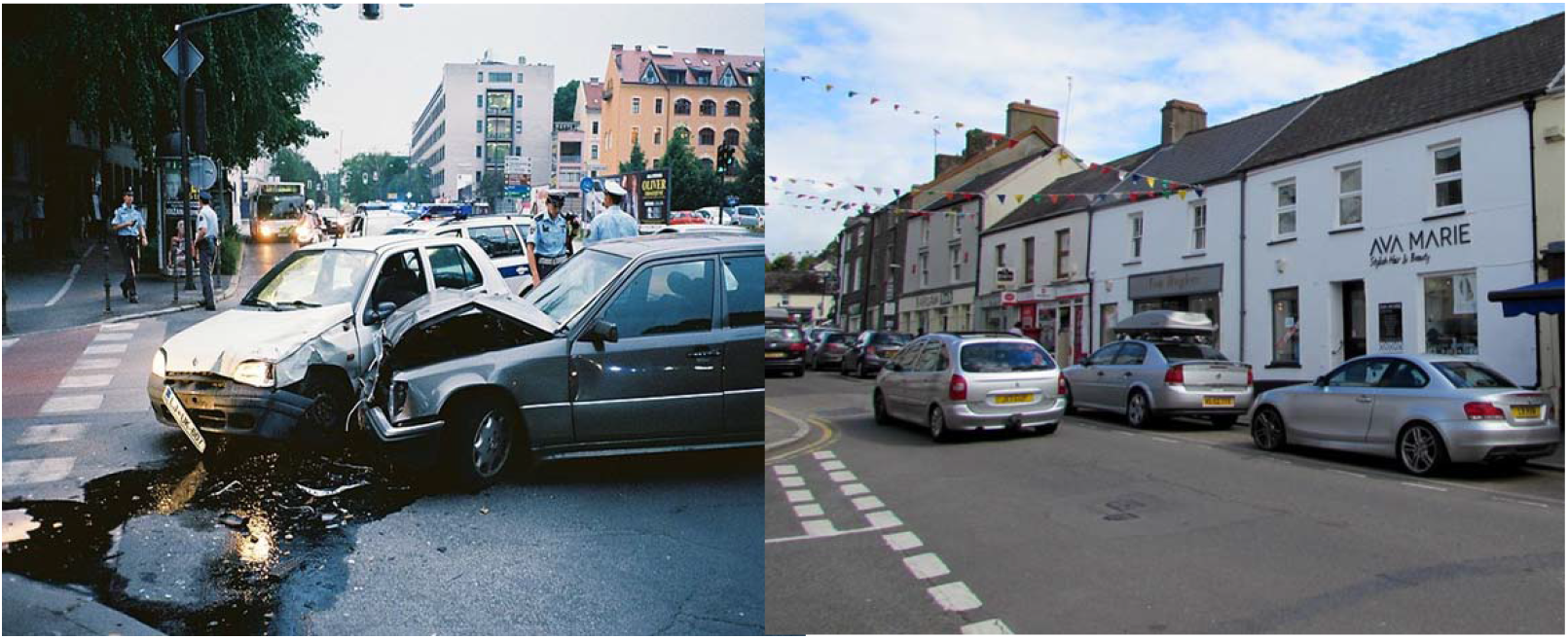
Example negative (emotionally arousing) and neutral stimuli. *Note. Left. Ljubljana car crash 2013 © Dino Kužnik (CC-BY-2.0) Right. High Street cars in Narberth © Jaggery (cc-by-sa/2.0*)

Partly because of the dominance of the modulated-consolidation model, much of the theory-building work on human emotional memory has been conducted at the level of the neurobiological mechanism (Cahill & McGaugh, 1998; Mather et al., 2015; McGaugh et al., 2000), contributing to an emphasis on consolidation processes and testing memory post-consolidation. It remains much less clear how emotional memory enhancement relates to the processes associated with the cognitive and computational neuroscience of memory. This is because factors that have traditionally interested human memory researchers, such as the level of processing at encoding, or the nature of the memory test, are questions at the algorithmic, rather than the implementation level. Talmi’s multi-factor mediation model (Talmi, 2013) discusses a number of these cognitive processes, but like all other existing models of enhanced emotional memory, her model, too, did not yield quantitative predictions. No existing framework has been implemented quantitatively in a formal model

The Modulated Consolidation model considers encoding processes only in the sense that the consolidation of experiences that were arousing during encoding would be modulated (Cahill, Gorski, & Le, 2003). The only existing model of the cognitive processes that underlie emotional memory, Arousal-Biased Competition (ABC) theory (Mather & Sutherland, 2011), and its neurobiological sister model, GANE, dissect the influence that arousal has on competitive attention allocation at encoding, and offers a rich account for the multiple routes that lead to the prioritisation of particular stimuli, and how priority and its downstream memory consequences are influenced by systemic arousal. These accounts are unique because they can account for memory for emotional experiences not only in delayed, but also in immediate tests. This is advantageous because memory for such experience is often enhanced in immediate tests, as measured by free recall, cued recall, and recollection (Dolcos, LaBar, & Cabeza, 2004; Kensinger & Corkin, 2004; Talmi, Schimmack, Paterson, & Moscovitch, 2007; see also Table 1 here).

Existing models of emotional memory enhancement do not consider the possibility that emotion could have additional influences during the retrieval stage. This limits their ability to relate to some of the age-old themes in the study of human memory, such as the nature of the test (recall, recognition), assumed retrieval process (memory search, recollection, familiarity), and the context of the test (similar or different to the encoding context). The relative neglect of retrieval mechanisms in these frameworks is incompatible with the mainstream human memory literature, where the consensus has long been that the conditions of retrieval often determine memory performance, as in the encoding specificity principle (Tulving & Thomson, 1973), transfer-appropriate processing (Graf & Ryan, 1990), and state-dependent memory (Smith & Vela, 2001) as well as their neural concomitant, instantiation (Danker & Anderson, 2010). Specifically in relation to emotion, the literature on mood congruency established that the retrieval context matters for emotional memory, with memory performance increasing when the mood at encoding and retrieval is consistent (Blaney, 1986). A study where participants encoded and retrieved stimuli with an emotional or a neutral context showed that this effect is not due solely to an emotional context at encoding or at retrieval, but to their match (Xie & Zhang, n.d.) Recent results also show that like neutral items (Cai et al., 2016), emotional items also have access to a drifting neural context (Rashid et al., 2016). Importantly, Rashid et al. showed that retrieving an emotional experience changes the way that a new experience is encoded, such that the two are functionally linked. It is reasonable, therefore, to expect that retrieval processes contribute to emotional memory.

### 1.2 The potential of retrieved context models in research of emotional memory

Memory is better when context states at encoding and at retrieval are more similar (Howard & Kahana, 2002a; Smith & Vela, 2001). Some have attributed the recency effect to such contextual similarity (Glenberg & Swanson, 1986; Neath, 1993). The term “brain state” has been employed recently (Deco & Kringelbach, 2017; Manns et al., 2007; Tambini, Rimmele, Phelps, & Davachi, 2017) to describe mental context that evolves and changes gradually with time (Bower, 1967; Estes, 1955). By moving from random fluctuations in mental context to context changes that are due to experiencing individual episodes or experimental stimuli, temporal context models have had much success in accounting for free recall dynamics, including recency effects (Howard & Kahana, 2002a; Sederberg et al., 2008). Famously, these models successfully account for the contiguity effect - the propensity to recall contiguously items that have been encoded close in time to each other. The contiguity effect is important because it contributes to our understanding of how memories are associated, and because it correlates with memory success (Sederberg, Miller, Howard, & Kahana, 2010). While the temporal context model based memory solely off of temporal associations (Howard & Kahana, 2002a), it has been clear for some time that semantic associations also contribute to retrieval success (Howard & Kahana, 2002b), and one of its updated versions, CMR (Polyn et al., 2009), formally included nontemporal dimensions of context.

CMR is a computational model that makes predictions about memory dynamics in free recall and recognition. Built on the notion that the content of a memory is intimately tied to its associated (internal) context, CMR assumes that each presented item is associated with a context state, and that the brain’s maintained state of context is updated by each presented or retrieved item. In this way, context changes slowly over time, as a recency-weighted sum of prior context states. During memory retrieval, the current context state is used to cue recall or recognition. CMR includes two dimensions of items’ non-temporal context, which are known to contribute to recall dynamics. Semantic context refers to the pre-experimental associations between studied items. Source context refers to experimental associations between studied items, resulting from specific encoding operations performed during the study of some items but not others.

In Polyn’s et al.’s first simulation this referred to the orienting task (size or animacy judgments). CMR also assumed that a switch between these two source contexts disrupts the temporal context, such that the temporal context states between two items separated by a task switch were less similar. The model was supported by evidence of clustering around the source context, concomitant with reduced reliance on temporal context (Polyn, Erlikhman, & Kahana, 2011; Polyn et al., 2009). Because similarity between the study and test contexts on any dimension can help retrieve a studied item, this generalisation enables CMR to explain the increased tendency to recall contiguously items that are similar to each other but were encoded far apart.

The fact that CMR considers the memory consequences of the similarity between encoded experiences beyond temporal context alone is important for the purpose of accounting for the effects of emotion on memory. If emotional items are represented as having a higher degree of similarity to one another on an ‘emotional context’ dimension, CMR should predict that participants will cluster their recall around the dimension of emotionality. There is, in fact, empirical evidence for emotional clustering of free recall. In one study participants encoded three pure lists of emotional, random-neutral and related-neutral items, and received a single, surprise final free recall test after all three lists were encoded (Talmi, Luk, McGarry, & Moscovitch, 2007). There was evidence for clustering around list type, and also evidence that such clustering had mnemonic consequences. In that study, temporal and emotional context effects were confounded because the three item categories were studied in three separate lists. Further research showed that emotional clustering appears also in free recall of mixed lists that contain both emotional and neutral items. For example, Barnacle et al. (Barnacle, Montaldi, Talmi, & Sommer, 2016) presented participants with mixed lists of emotional and neutral pictures which were equally semantically related, and found clustering around the emotional/neutral category; the degree of clustering predicted memory performance. Long and colleagues (Long, Danoff, & Kahana, 2015) presented participants with mixed lists of words, some of which were negative and some positive, and found that participants tended to retrieve positive items after other positive items, and similarly for negative and neutral items. Emotional clustering was observed even when pre-experimental semantic associations between words (obtained through latent semantic analysis scores) were taken into account. In another study, the transition probability from pleasant items to pleasant items, and from unpleasant items to other unpleasant items, was higher than the transition from neutral to neutral items (Siddiqui & Unsworth, 2011). These findings suggest that participants retrieve the emotional context of recalled items, which updates the test context and becomes part of the next retrieval cue.

### 1.3 The emotional Context-Maintenance and Retrieval (eCMR) model

Here we generalize the CMR framework to examine hypotheses regarding the emotional modulation of memory. We take advantage of CMR’s representations along multiple stimulus dimensions to incorporate emotionality of presented items. In this section, we consider systematically which aspect of CMR may be influenced by emotion. Our extended model, eCMR, is described formally in section 2.2 and schematically in FIGURE 2. For expository purposes, we introduce the two changes we propose as nested model variants. The *category-only variant* represents certain items as belonging to an emotional category, and others as belonging to a neutral category. The *attention-category* variant (depicted in Figure 2) adds a modulatory effect of emotional arousal on simulated attention. We show that the attention-category variant, where emotion is operationalised in the model through two different mechanisms, is better than able to capture a number of key emotional memory phenomena.

**FIGURE 2:**
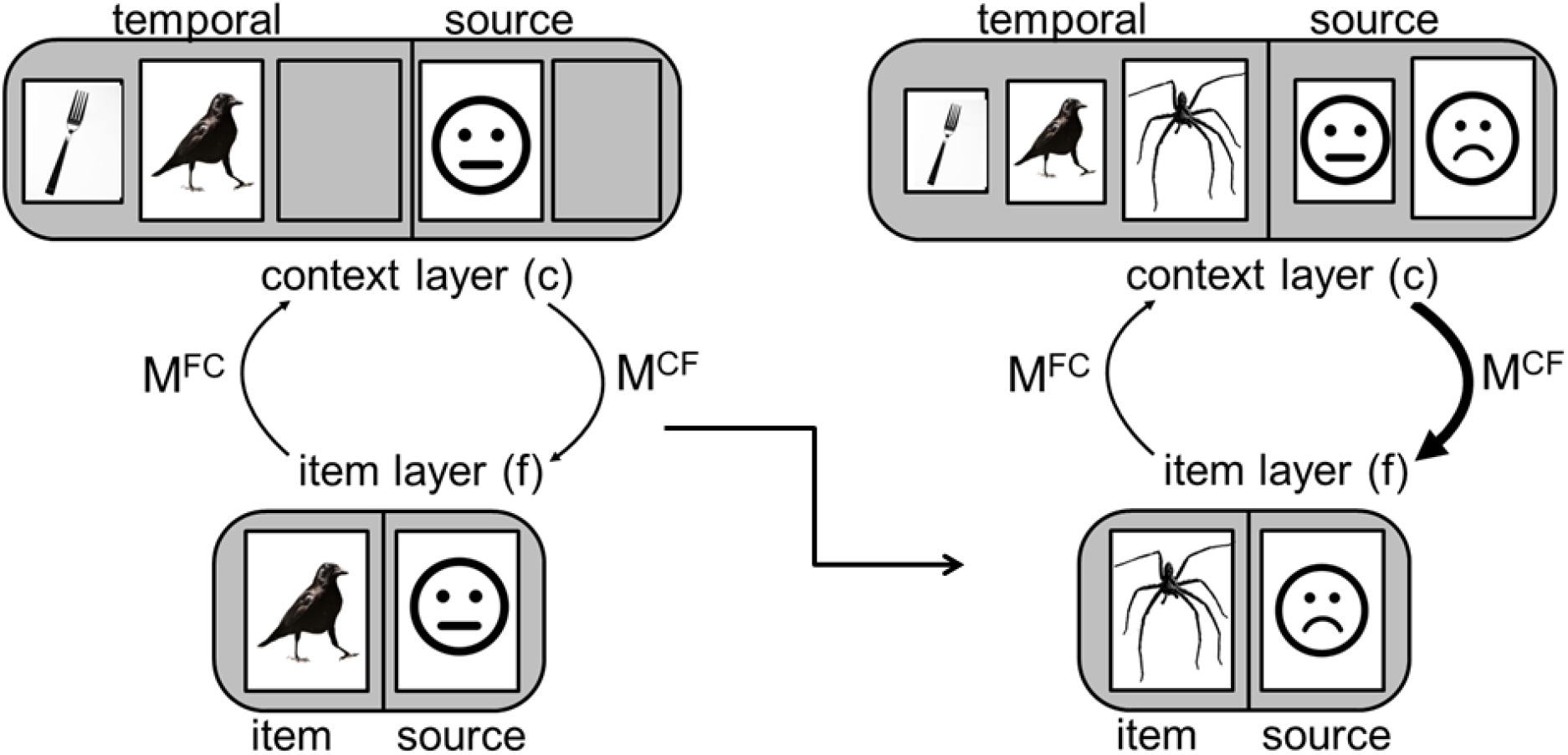
A schematic of the encoding of two items in eCMR. **Left.** A new item, bird, is encoded in the feature layer, with a temporal feature reflecting its second position in the list, and the emotional feature ‘neutral’. It is associated with the context layer, where it dominates the temporal and emotional context sub-regions over the previous item (fork, also neutral). **Right.** A third item is encoded into the feature layer, with a temporal feature reflecting its third position in the list, and the emotional feature ‘arousing’. These update the context layer, dominating over the previous item. The context resembles the most recently-encoding item and facilitates the recall of the spider over the bird. The thicker arrow from the context to the item layer represents the stronger association between these layers when the item has an emotional value in the attention-category variant of eCMR.

#### 1.3.1 The category-only variant of eCMR

This variant introduces the way emotion is built into the structure of eCMR as a source context. In CMR (Polyn et al., 2009) each item is represented with three types of features: item, semantic, and source. The item features are typically uniquely set for each item. Semantic features of the items are incorporated into the association matrix between temporal context and item features, under the assumption that semantically related items co-occur in similar contexts over time (Rao & Howard, 2008). Each item also has a ‘source’ feature corresponding to aspects of the stimulus that are common to many items in the study set, such as the orienting task that was associated with each item during encoding. Polyn and colleagues recognized that source features could be internal – unique operations that participants may engage in during the encoding of certain stimulus types. When items are encoded their temporal and source contexts are updated, resulting in increased similarity between them. When a particular item is recalled, it promotes the recall of items with similar semantic, temporal, and source context dimensions.

Consider the emotional items that are prevalent in emotional memory experiments, such as a picture depicting a car crash (FIGURE 1). In the *category-only variant* of eCMR the ‘source’ item feature implements its emotionality, such that it defines certain items as ‘emotional’ and others as ‘neutral’. Graded coding is entirely feasible but for the purpose of the simulations we present here, we defined items in a binary way as ‘emotional’ or ‘neutral’. Just as in CMR, where items that share source context promote each other’s recall, in eCMR items that share emotional context will also promote each other’s recall.

The temporal and semantic features are represented in exactly the same way in eCMR as in CMR, but the strength of semantic associations operationalise both the semantic similarity between items, based on their shared conceptual and thematic relationship, and their emotional similarity. By definition, emotional stimuli belong to the same conceptual category (e.g. ‘negative experiences’). Emotional stimuli are often also thematically similar; for example, the words ‘bomb’ and ‘starvation’ will co-occur within a story of war. Other factors that may increase the thematic similarity between emotional items is that they may be construed as socially more ‘distant’ (Trope & Liberman, 2003), or share affordances (Greene, Baldassano, Esteva, Beck, & Fei-Fei, 2016). Emotional stimuli may also be similar because they trigger the same feelings, embodied in a myriad of peripheral physiological processes (Bradley, Lang, & Cuthbert, 1993). The documented increased similarity between emotional stimuli compared to randomly-selected neutral ones has memory consequences, explaining part of the recall advantage of emotional items (Talmi, Luk, et al., 2007; Talmi & Moscovitch, 2004). It is important to note that some experiments go to great lengths to equate the pre-experimental similarity of emotional and neutral stimuli (Kensinger, 2009b; Schmidt & Saari, 2007; Sommer, Gläscher, Moritz, & Büchel, 2008), including the key experiment we simulate in section 2 (Talmi et al., 2007). However, even when pre-experimental similarity is controlled, emotional items will still end up more similar to each other in eCMR because during encoding they will be bound to a shared emotional context.

Lastly, a critical aspect of CMR is the assumption that the temporal context is disrupted whenever the orienting task changes during encoding (Polyn et al., 2009). While the task Polyn and colleagues used is akin to task-switching paradigms, mixed lists of emotional and neutral items may more closely resemble cue-shift paradigms, where the task remains the same, but the cue that lets participants know which task to execute varies. Just as an animacy-judgment task could be cued by both a circle and a square, the operations that emotional items trigger are cued by items with different content. Importantly, both task-switches and cue-switches slow reaction times in the post-shift trials (Schneider & Logan, 2006), a result that may indicate that both type of switches could be modelled as disrupting the temporal context. As we will see, how much the temporal context is disrupted when emotional and neutral items alternate substantially influences eCMR’s predictions. In keeping with CMR, in section 2 we assume that such disruption occurs, albeit subtly, while in section 5 we examined the impact of a substantial disruption. In summary, the *category-only variant* of eCMR in Simulation 1 is very similar to CMR but uses emotionality as an internal source feature of items and a lower level of temporal context disruption with a change in task.

#### 1.3.2 The attention-category variant of eCMR

This variant subsumes the category-only variant but allows emotion to modulate attention during encoding. Emotionally-arousal enhances attentional selection, modulation and vigilance (Golomb, Turk-Browne, & Chun, 2010). For example, autonomic arousal or visual attention to objects that have been previously presented on emotional backgrounds is enhanced even when those emotional backgrounds are no longer there (Ventura-Bort et al., 2016). Allocation of attention to emotional stimuli is driven by the amygdala and results in enhanced sensory processing downstream, a mechanism which is distinct from (direct) stimulus-driven effects or strategic top-down effects (Pourtois, Schettino, & Vuilleumier, 2013). The mechanism involves amygdala-dependent activation of the locus coeruleus, where noradrenergic neurons increase the neural gain throughout the brain through their wide-ranging projections (Mather et al., 2015). Based on the consensus in the literature eCMR assumes that stimuli with an ‘emotional’ item feature attract attention preferentially, although of course, in reality there is variability in the degree of attention allocated to specific emotional stimuli.

In retrieved context models such as CMR, CMR2 and TCM-A (Lohnas et al., 2015; Polyn et al., 2009; Sederberg et al., 2008), increased attention to early stimuli in the study list has been implemented by strengthening the associations between items and their temporal contexts. This is accomplished by modulating the step-size or learning rate parameter on encoding these items, itself a standard mechanism to capture competitive attentional effects in associative learning theories more generally (Dayan, Kakade, & Montague, 2000; Pearce & Hall, 1980). Empirically, attention at encoding improves recall; for example, late positive potential at study, an ERP sensitive to stimulus-driven attention, covaries with memory success (Chen, Lithgow, Hemmerich, & Caplan, 2014). In retrieved context models the tighter binding of items to their context likewise increases the competitiveness of items during retrieval.

The implementation of attention in eCMR as increased strength of item-context associations corresponds to evidence that emotional stimuli are more tightly associated to their context. According to the priority-binding hypothesis (MacKay et al., 2004) emotional items are bound more strongly to their presentation context than neutral items. This assertion is evident in that emotional items escape the attentional blink more readily – namely, they are more likely to be reported when presented in a rapid serial visual stream close to a target stimulus – because they are bound better to their encoding context (Anderson, 2005; Mackay, Hadley, & Schwartz, 2005). Similarly, in the object-based framework attention is allocated preferentially to features perceived as belonging to the same emotional object, driving increased binding of emotional object features (Mather, 2007). The object-based framework is supported by evidence for increased memory for the source of within-object features, such as the font colour and screen location of taboo words (MacKay & Ahmetzanov, 2005). In summary, the *attention-category* variant of eCMR in Simulation 2 encapsulates the *category-only variant* but also makes one additional assumption - that emotional items attract more attention, operationalised as increased strength of their binding to their encoding context. After we present the model formally we discuss the different ways that increased binding to the context could be implemented.

To preview how this eCMR variant works, consider that emotional stimuli are attended preferentially when they are presented. Therefore, they are more strongly bound to their context at encoding, compared to neutral items, and – when the model simulates recall of mixed lists – they are more strongly associated to the context at the beginning of the memory test, increasing the likelihood that recall would begin with an emotional, rather than a neutral item. Encoding emotional items renders the source context of an immediate test more emotional than it would be after the encoding of neutral items; the number of emotional items in the list, particularly at the late serial positions, will influence the emotionality of the test context. An emotional context will facilitate recall of emotional items. Likewise, recalling an emotional item promotes the recall of another emotional item, both because they share their source context, and because during encoding, the switch to an emotional source disrupted the temporal context, reducing the likelihood that previously- or subsequently-encoded neutral items would be retrieved on the basis of temporal contiguity. Although many of the experiments simulated here control for semantic relatedness, another reason that emotional items promote each other’s recall is that they are often also more closely related to each other semantically than other items. Each recall of an emotional item additionally updates the temporal context, moving it yet further away from the temporal context that pertained during the encoding of neutral items, and therefore further hindering their recall. This variant of eCMR thus describes emotional memory enhancement as multiply determined, in keeping with the multiple factors previously posited to contribute to this effect (Talmi, 2013).

### 1.4 Overview of aims and objectives

Thus far we have delineated, in broad terms, the limitation of previous models of emotion-enhanced memory and the promise of an extension of CMR – eCMR – to account for critical effects of emotion on episodic memory. Our overall aim in this paper is to illustrate how this can be achieved on a qualitative level for all effects, rather than fit the nuances of any particular data set. We chose this approach because a more precise fit would most likely require a rigorous parameter search for each data set. With different parameter values for each data set, this might obfuscate whether the model can only capture each of the effects within restricted ranges of specific parameter values, or can capture all effects simultaneously. By comparing model predictions on a qualitative level we could keep constant more model parameter values. In fact, we set as many parameters possible based on a previous algorithmic search for best-fit parameters in free recall (Polyn et al. 2009). For other parameters with changed values, we nonetheless kept most values constant across simulations. Within each section parameter values only change to simulate the study-test retention interval and the increased surprise occasioned by the presentation of an oddball in the list (Tables 1–2). Across sections we only varied the relative reliance on semantic associations, because section 2 used more cohesive stimuli.

**TABLE 1.**
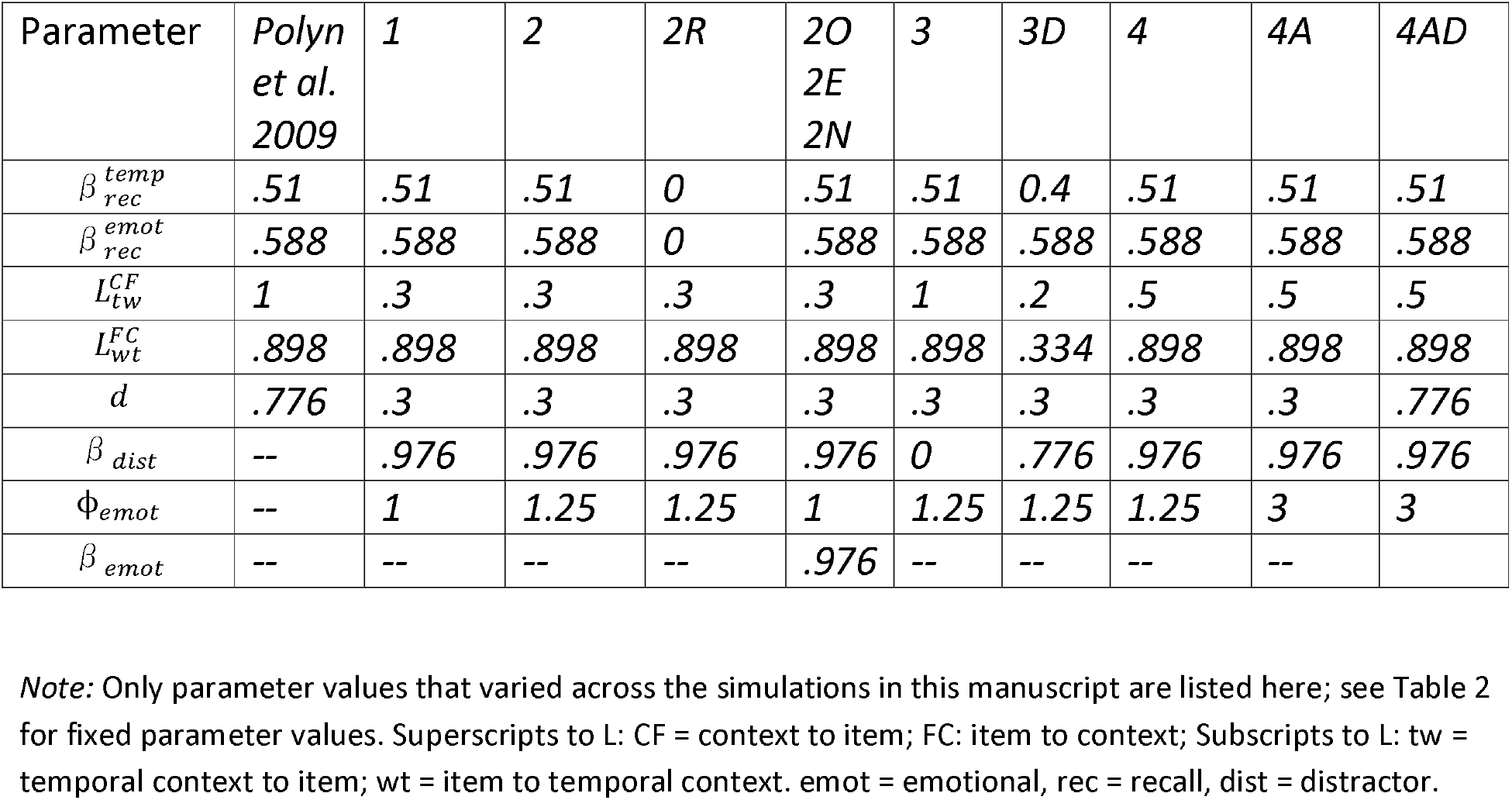
Variable parameter values in Simulations 1-5.

| Parameter | *Polyn et al. 2009* | 1 | 2 | 2R | 2O<br>2E<br>2N | 3 | 3D | 4 | 4A | 4AD |
| --- | --- | --- | --- | --- | --- | --- | --- | --- | --- | --- |
| $\beta_{rec}^{temp}$ | .51 | .51 | .51 | 0 | .51 | .51 | 0.4 | .51 | .51 | .51 |
| $\beta_{rec}^{emot}$ | .588 | .588 | .588 | 0 | .588 | .588 | .588 | .588 | .588 | .588 |
| $L_{tw}^{CF}$ | 1 | .3 | .3 | .3 | .3 | 1 | .2 | .5 | .5 | .5 |
| $L_{wt}^{FC}$ | .898 | .898 | .898 | .898 | .898 | .898 | .334 | .898 | .898 | .898 |
| $d$ | .776 | .3 | .3 | .3 | .3 | .3 | .3 | .3 | .3 | .776 |
| $\beta_{dist}$ | -- | .976 | .976 | .976 | .976 | 0 | .776 | .976 | .976 | .976 |
| $\phi_{emot}$ | -- | 1 | 1.25 | 1.25 | 1 | 1.25 | 1.25 | 1.25 | 3 | 3 |
| $\beta_{emot}$ | -- | -- | -- | -- | .976 | -- | -- | -- | -- | |

*Note:* Only parameter values that varied across the simulations in this manuscript are listed here; see Table 2 for fixed parameter values. Superscripts to L: CF = context to item; FC: item to context; Subscripts to L: tw = temporal context to item; wt = item to temporal context. emot = emotional, rec = recall, dist = distractor.

**TABLE 2.**
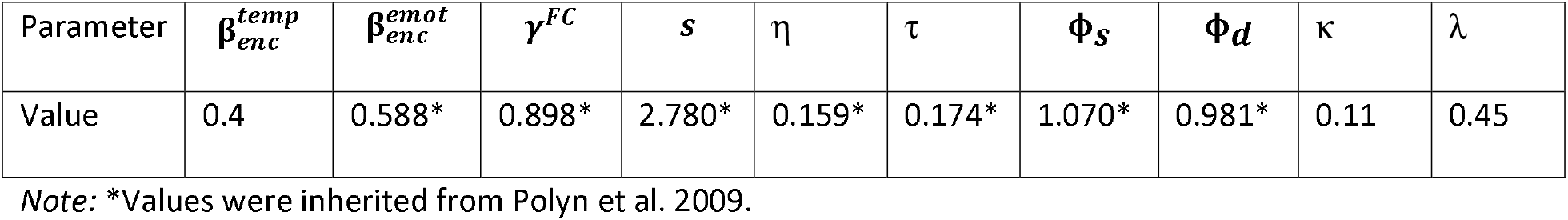
Fixed parameter values in simulations.

| Parameter | $\beta_{enc}^{temp}$ | $\beta_{enc}^{emot}$ | $\gamma^{FC}$ | $s$ | $\eta$ | $\tau$ | $\phi_s$ | $\phi_d$ | $\kappa$ | $\lambda$ |
| --- | --- | --- | --- | --- | --- | --- | --- | --- | --- | --- |
| Value | 0.4 | 0.588* | 0.898* | 2.780* | 0.159* | 0.174* | 1.070* | 0.981* | 0.11 | 0.45 |

*Note:* \*Values were inherited from Polyn et al. 2009.

In section 2 we use the *category-only* and *attention-category* variants of eCMR to simulate the emotional list composition effect: the advantage of emotional over neutral items in the recall of mixed lists, and their diminished advantage in the recall of pure lists. While this is a relatively simple effect, described by four data points that correspond to average free recall, it has so far evaded explanation by existing models of emotional memory. We show that the *attention-category* variant captures this effect qualitatively, while the *category-only* variant failed to do so. By probing the retrieval dynamics in eCMR we show how the model helps reveal the multiple mechanisms that give rise to this effect and render it so robust. In section 3 we use the *attention-category* variant of eCMR to compare immediate and delayed recall of mixed lists, where the pattern of results reported in the literature on delayed emotional memory is less familiar in the literature of memory modelling. In Section 4 we examine a particularly challenging effect for eCMR, the emotional oddball effect. The attention-category variant of eCMR can capture the effect by assuming that more attention is allocated to the oddball and that it disrupts the temporal context of encoding more than a similar emotional item in mixed lists, where such items are more frequent. With these assumptions in place eCMR captures key empirical results.

## 2. eCMR captures the emotional list composition effect

The list-composition effect is an umbrella term that refers to an interaction between the way to-be-remembered items are processed during encoding and the global composition of the encoding list (McDaniel & Bugg, 2008). This manipulation could be executed by selecting atypical items with unique attributes that attract special encoding processes, for example those that are unusual, complex, or bizarre; or it could be due to experimental instructions to process a subset of items in an atypical way, for example by enacting or generating a subset of items while others are silently read. In tests of free recall, atypical items are recalled better when they are encoded in the same list as the ‘standard’ items, but their advantage is minimised or even eliminated when each item type is encoded separately, in a pure list (McDaniel & Bugg, 2008). In recognition memory tests memory for atypical items is better than memory for standard items (McDaniel & Bugg, 2008).

The list composition effect is related to the list-strength effect (Ratcliff, Clark, & Shiffrin, 1990) – the finding that spaced repetition of some of the items gives them a larger free-recall advantage when they are presented in a mixed list with non-repeated items, compared to a situation where repeated and nonrepeated items are presented in pure lists. Spaced repetition is thought to ‘strengthen’ the items, and is therefore akin to the manipulations mentioned above that render them ‘atypical’. The effects resemble each other also in that the list-strength effect also disappears in recognition memory tests (Shiffrin & Steyvers, 1997). Yet the effects of spaced repetition may not be the same as the effect of unusualness. Atypical items are thought to be recalled better in mixed lists because they are better attended and elaborated (McDaniel & Bugg, 2008), while longer or deeper processing of standard items does not give rise to the list-strength effect (Malmberg & Shiffrin, 2005).

Because emotional stimuli have unique attributes and attract unique processing operations, it is unsurprising that emotion yields an *emotional* list-composition effect in free recall. Recalling mixed lists operationalises telling a friend about a day that included some emotionally-arousing aspects: monthly targets were tabulated, the lunch was terrible, budget meetings were held, a close colleague announced they were leaving. Recalling pure lists operationalise telling someone about a more difficult day, with many emotionally arousing aspects: a child banged their knee, we got a ticket while rushing to school, and had a migraine the rest of the day. While the emotional stimuli are recalled robustly more than neutral ones when encoded in mixed lists, this memory advantage is smaller, and sometimes eradicated, in pure lists. Table 3 lists some demonstrations of that effect in the literature; note that the pattern is clearest when emotional stimuli are highly arousing (i.e. pictures or taboo words), and when the semantic relatedness of stimuli is equated. The influence of the retrieval test on the emotional list-composition effect should be studied further.

**TABLE 3.** The emotional list composition effect.

|  | Mixed<br>Emotional | Neutral | Pure<br>Emotional | Neutral |
| --- | --- | --- | --- | --- |
| (Barnacle et al., 2016)<sup>R</sup> | 59 | 58 | 64 | 48 |
| (Barnacle, 2015)(Chapter 6)<sup>R</sup> | 61 | 48 | 53 | 56 |
| (Barnacle, Tsivilis, Schaefer, & Talmi, 2017)<sup>R</sup> | 63 | 41 | 57 | 53 |
| (Dewhurst & Parry, 2000)(Exp 1 and 2)<sup>ac</sup> | 49 | 48 | 55 | 35 |
| (Hadley & MacKay, 2006)(Exp 2)<sup>b</sup> | 45 | 36 | 44 | 45 |
| (Sommer et al., 2008)<sup>P</sup> |  |  | 42 | 43 |
| (Talmi & McGarry, 2012)(Exp1)<sup>R</sup> | 56 | 31 | 56 | 42 |
| (Talmi & Moscovitch, 2004)(Exp 2B and 3)<sup>R</sup> | 27 | 33 | 22 | 36 |
| (Talmi, Luk, et al., 2007 (Exp 2A and 2B)<sup>R</sup> | 58 | 47 | 57 | 56 |
| (Talmi, Luk, et al., 2007 (Exp 4)<sup>R</sup> |  |  | 61 | 67 |
| (Talmi, Luk, et al., 2007)(Exp 1) | 72 | 57 | 71 | 66 |
| (Watts, Buratto, Brotherhood, Barnacle, & Schaefer, 2014) | 37 | 25 | 35 | 30 |

*Note. Other than when indicated, all data refers to percentage recall in free recall tests that used picture stimuli. Data in the table was extracted from the figures when the averages were not reported in the text.*<sup>a</sup>
*Emotional word stimuli.*<sup>b</sup> *taboo word stimuli.*<sup>c</sup> *Corrected “Remember” responses.*<sup>R</sup> *stimuli were controlled for semantic relatedness.*<sup>P</sup> *Arousing and non-arousing items, data kindly provided by the author.*

The emotional list-composition effect is outside the scope of the modulated-consolidation model and the emotional binding account, because it is obtained in immediate memory tests. It is also difficult to account for the effect using GANE or ABC theory because of evidence that emotional items receive extra attention in both mixed and pure lists (Barnacle & Talmi, 2016). For example, when participants viewed stimuli under divided attention their performance on the concurrent auditory choice reaction time task was poorer when stimuli were emotional in both pure and mixed lists (Talmi & McGarry, 2012). In another study EEG was recorded during encoding. Electrophysiological markers of attention and working memory were increased when participants viewed emotional stimuli in both pure and mixed lists (Barnacle, Tsivilis, Schaefer, & Talmi, 2018). In these studies list composition did not modulate attention towards emotional stimuli. Indeed, this appears to be a common feature across list composition manipulations (reviewed in McDaniel & Bugg, 2008). Furthermore, in a study where we scanned participants with fMRI during encoding in the emotional list-composition paradigm we found no evidence for reduced attention to neutral stimuli or increased attention to emotional stimuli in mixed, compared to pure lists (Barnacle et al., 2016). These empirical findings motivated us to assume, in Simulations 1-2, that increases in attention to emotional compared to neutral items were equivalent in magnitude across pure and mixed lists. Taken together, because emotional stimuli are atypical, and because they are attended and elaborated more than neutral stimuli; and because atypical stimuli produce list-composition effects even when they are not emotional, all list composition effects may share some of their mechanisms. By contrast, the emotional list-composition effect is probably less related to the list-strength effect, which depends on spaced repetition but is unaffected by elaboration.

Below we describe two ways of operationalising the impact of emotion on CMR based on the considerations discussed thus far. The test for each model variant is its ability to mimic qualitatively the empirical pattern of results depicted in **Error! Reference source not found.,** rather than reproduce the numbers themselves. Specifically, we were looking to capture the interaction between emotion and list composition, where emotion enhances recall in mixed but not pure lists. After we describe the simulations of average recall we describe recall dynamics, and then additional empirical data that supports the conclusion that the emotional list composition effect depends on multiple aspects of encoding and retrieval dynamics.

### 2.1 A description of the empirical data from Talmi et al., 2007

We simulated the average recall data from the emotional and related-neutral conditions in Talmi et al., 2007, Experiment 2. Details of the methods of that experiment are presented in **APPENDIX 1**. The semantic coherence of the stimuli used in that experiment was matched by selecting neutral items that depicted domestic scenes, and selecting emotional and neutral stimuli matched on their average inter-relatedness based on a separate rating study.

The results of this experiment, which are depicted in **Error! Reference source not found.,** produced the pattern we refer to in this paper as the emotional list-composition effect. Emotional stimuli were recalled better than neutral ones in the mixed list condition, but their advantage was diminished, and here nonsignificant, in the pure-list condition. The pure-list condition, which is rarely employed in research on emotional memory, offers a unique insight about the mixed-list results. Essentially, the pure list condition could be considered a control condition, against which the mixed list condition could be interpreted. Described in this way, the data suggest that the emotional-memory advantage in mixed lists stems entirely from a decrease in memory for neutral items in the mixed-list condition, compared to the pure-list condition, rather than from an increase in memory for emotional items in mixed lists. Compared to pure lists, across experiments we consistently find a decrease in memory for neutral items presented in mixed lists, but memory for emotional items in mixed lists is only increased some of the time, as in the second experiment in **Error! Reference source not found.** (Barnacle et al., 2018), a variability that awaits further research.

To fully understand the interaction between attention to emotion at encoding and the retrieval machinery of eCMR we analysed the types of transitions the model made between successive recalls in the empirical data. Power was slightly higher in Barnacle et al.’s (2018) study because there were four lists in each condition there compared to one list in Talmi et al.’s(2007) study. First, we examined the proportion of transitions made based on temporal associations in pure lists, and how those were affected by category membership (emotional/neutral). **Error! Reference source not found.** depicts contiguity effects in the recall of pure lists, plotting the probability of recalling each item as a function of the lag from the currently-recalled item at lag=0 (Howard & Kahana, 1999). It is clear that these curves are very noisy; the number of lists they are based on is very small compared to other studies of recall dynamics in the literature. The plots suggest that the empirical lag-CRP curves have shallow slopes, indicative of limited reliance on temporal clustering; and an apparent trend for increased recall of emotional items at lag+1. To increase power we collapsed the across serial positions and computed temporal clustering scores (Polyn et al., 2011, 2009). For each recall transition, we considered the absolute lag of that transition against the distribution of absolute lags from not-yet-recalled items, assigning it a score from 0 to 1, where 1 reflects that the transition was the smallest absolute lag possible. Thus, overall higher temporal clustering scores reflect recall organisation with greater influence from temporal associations, and 0.5 the baseline level expected by chance (TABLE 4.). Temporal clustering was greater than chance in the more powerful Barnacle et al. dataset (emotional lists: *t*(24)=3.27, p<0.01; neutral lists: *t*(24)=2.15, p<0.05) but not in the Talmi et al. dataset (emotional lists: *t*(23)=1. 11, p=.27; neutral lists: *t*(23)=1.91, p=.06). In neither was there a significant difference between conditions (Talmi et al. t<1, Barnacle et al., t<1). The low clustering scores again suggest limited reliance on the temporal context during recall in that experiment. Two aspects of the task might have contributed to this. First, each experimental list was preceded and followed by two buffers, and a distractor task was interpolated between study and test, removing the majority of primacy and recency influences. Second, the majority of experimental stimuli were related semantically, increasing reliance on semantic and emotional dimensions of context compared to the temporal context. Our choice of model parameters was informed by the relative contributions of temporal organization and emotional organization to recall dynamics. Given the noisiness of the contiguity scores we did not at present attempt to simulate the increased recall of emotional items at lag+1, and note this may be an area where further model development is warranted, with a more powerful dataset. We comment on this issue in further detail in section 2.5.

**TABLE 4.**
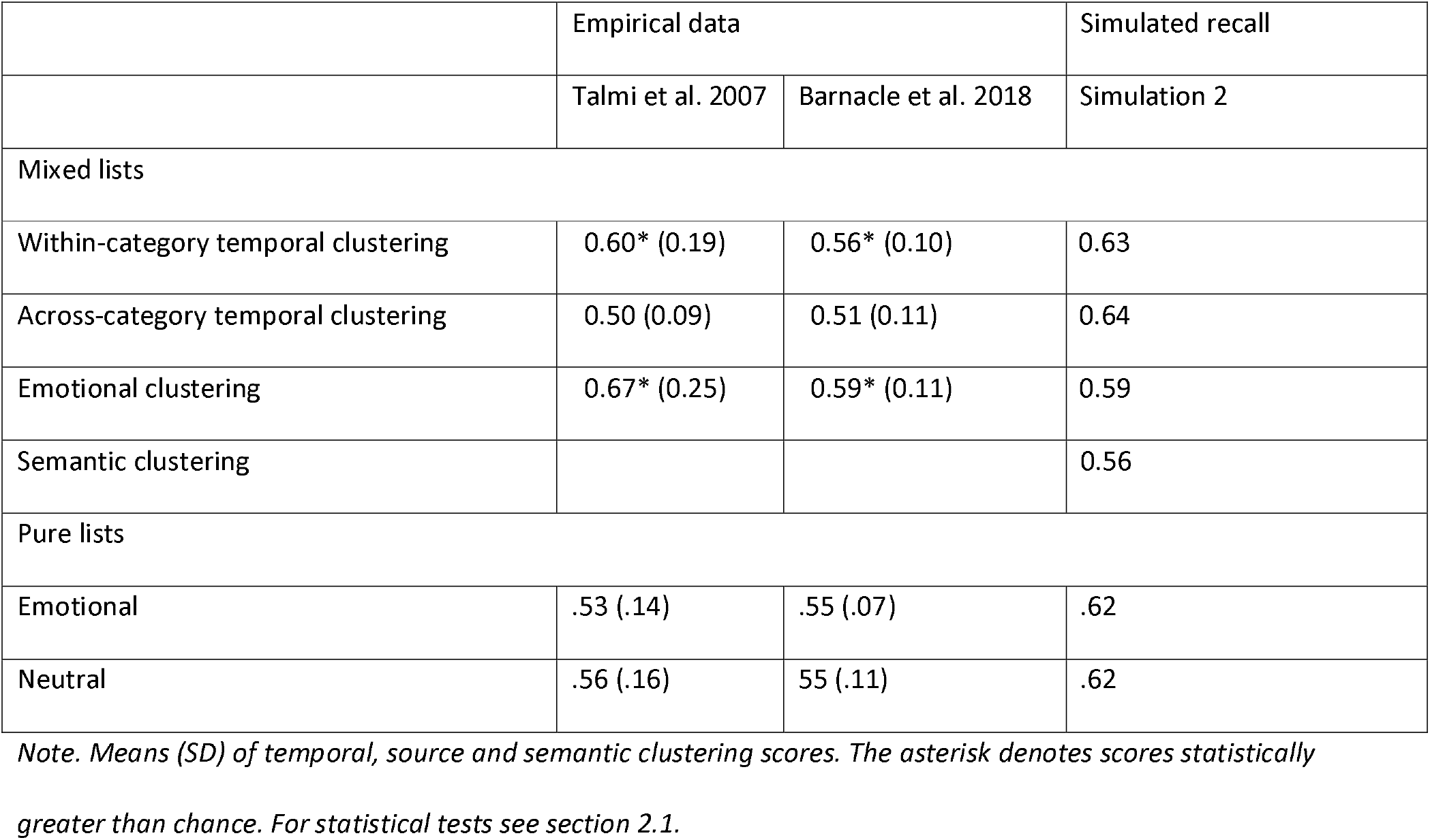
Empirical and simulated recall transitions.

|  | Empirical data |  | Simulated recall |
| --- | --- | --- | --- |
|  | Talmi et al. 2007 | Barnacle et al. 2018 | Simulation 2 |
| Mixed lists |  |  |  |
| Within-category temporal clustering | 0.60* (0.19) | 0.56* (0.10) | 0.63 |
| Across-category temporal clustering | 0.50 (0.09) | 0.51 (0.11) | 0.64 |
| Emotional clustering | 0.67* (0.25) | 0.59* (0.11) | 0.59 |
| Semantic clustering |  |  | 0.56 |
| Pure lists |  |  |  |
| Emotional | .53 (.14) | .55 (.07) | .62 |
| Neutral | .56 (.16) | .55 (.11) | .62 |

*Note. Means (SD) of temporal, source and semantic clustering scores. The asterisk denotes scores statistically greater than chance. For statistical tests see section 2.1.*

Next, we examined transitions in the recall of mixed lists (TABLE 4.). Transitions that were based on temporal contiguity within the same category, namely, the tendency to retrieve an emotional (neutral) item that was encoded close in time to another emotional (neutral) item, was greater than chance (Talmi et al. *t*(22)= 2.49, *p*=.02; Barnacle et al. *t*(23)=2.72, *p*=.01). This was not the case for temporal clustering between categories, which was at chance (Talmi et al. *t<1*, Barnacle et al., *t<1*). We also examined the emotional clustering score, defined as the proportion of transitions made to the same emotional category out of all transitions. Thus, recall of an item from either emotional state (neutral or emotional) will support recall of items from the same emotional state, regardless of any temporal contiguity. Transitions based on shared emotional context were, again, greater than chance (Talmi et al. *t*(23)= 3.31, *p*<.01, Barnacle et al. *t*(23)=3.77, *p*<.001).

**FIGURE 3.**
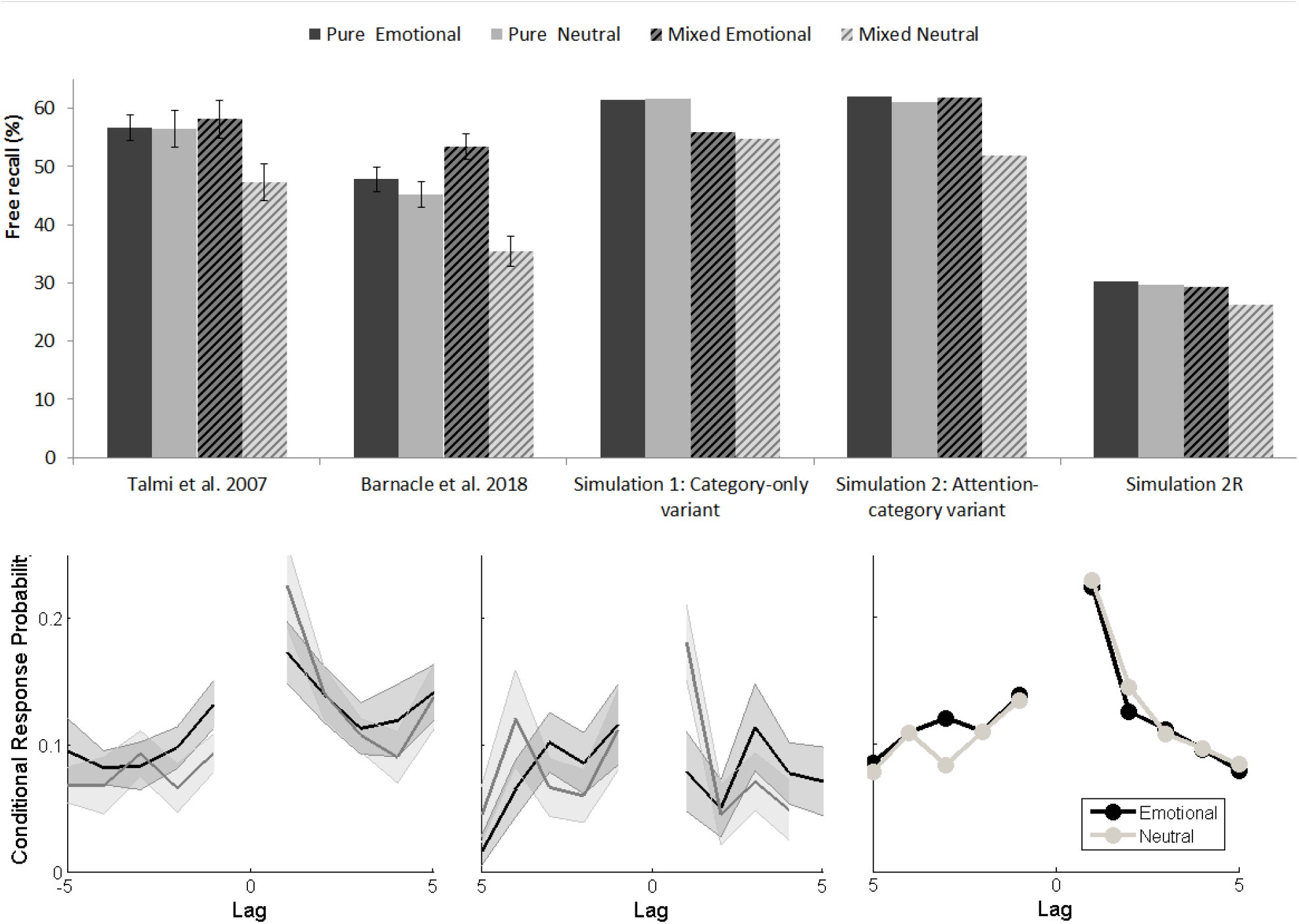
Recall data from Talmi et al., 2007, Experiment 2 (left) and the results of Simulations 1-2. ***Top.** Average recall data. The attention-category (Simulation 2) variant of eCMR captured the emotional list composition effect, where emotion enhances memory in mixed but not pure lists. The category-only variant (Simulation 1) failed to do so. Error bars refer to the standard error of the mean*. ***Bottom.** Contiguity effects in the recall of pure lists. **Left:** in Talmi et al. (2007). **Middle:** in the recall of pure li and Barnacle et al. (2018). **Right:** Predicted contiguity effects in the attention-category variant of eCMR (Simulation 2). Black: emotional lists*. *Grey: neutral lists. The shaded thickness represents standard error*.

### 2.2 A formal description of eCMR

We begin by describing the assumptions of eCMR that follow directly from those of CMR, before turning to those unique to a model that incorporates the emotional aspect of memory. In eCMR, each studied item *i* has an associated feature vector **f**_*i*_ and context vector **c**_*i*_ which interact through the association matrices *M^FC^* and *M^CF^*. When an item is presented to the model, this activates its feature vector **f**_*i*_. This vector is a concatenation of item features 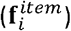 and emotional features 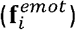, analogous to the definition of item and source features in CMR. For simplicity, each temporal context and emotional context sub-region of **f**_*i*_ has a localist, orthonormal representation.

This feature vector then creates an input to context. This input to context, 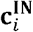, is defined as:

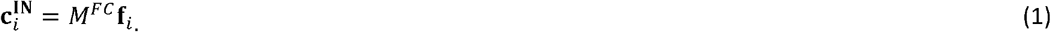

Like the item vector, the context vector is a concatenation of temporal and emotional representations, and each of the temporal and emotional contexts is updated separately and normalized to have unit length. Next, this input to context, 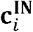, is used to updated the current context state, **c**_*i*_:

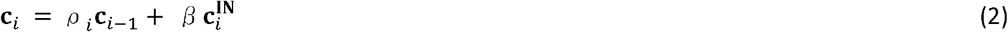

where *β* defines how much context is updated for each presented item, and takes on a separate value for the temporal and emotional sub-regions (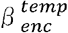 and 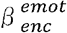, respectively). This arrangement allows the emotional context to drift at a different rate than the temporal context. Given that physiological arousal can give rise to slow systemic effects, this was seen as a desirable property. The *β* parameters are fixed to the same value for each presented item, *ρ_i_* is determined separately for each sub-region to normalize the level of contextual activation to have unit length (see Howard and Kahana 2002 for a more detailed discussion of the importance of this step):

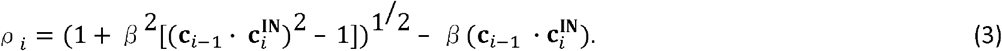

As each item is presented, the associative matrices (*M^FC^* and *M^CF^*) are updated according to a standard Hebbian learning rule, such that:

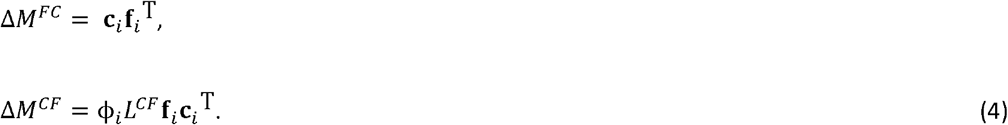

For early list items, eCMR assumes a primacy gradient of attention such that the change in *M^CF^* is scaled by ϕ_*i*_, which is greatest for early list items and decreases exponentially to an asymptotic value over the course of list presentation:

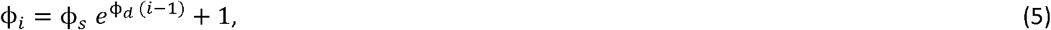

where ϕ_*s*_ and ϕ_*d*_ are model parameters. *L^CF^* in Equation 4 is a matrix that allows CMR to scale the magnitude of source associations relative to temporal associations. *L^CF^* has four sub-regions, corresponding to each of the 4 possible association types:

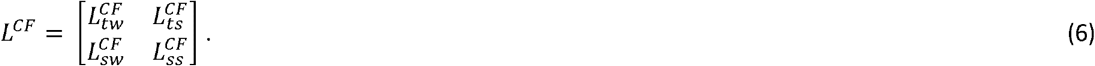

In the subscript, the first term refers to the context type (temporal (*t*) or source (*s*), i.e. emotional), and the second term refers to the feature type (again, temporal (*w*) or source). For simplicity, only the temporal feature terms are set to non-zero values: the emotional context to temporal feature associations, 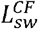, is a model parameter; the temporal context to item feature associations, 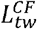, was fixed at 1 for Polyn et al., 2009, but we consider other values for this parameter, as described below.

In a similar way, *M^FC^* also has 4 sub-components (i.e. those starting with *w*), but again only the temporal feature terms are set to non-zero values:

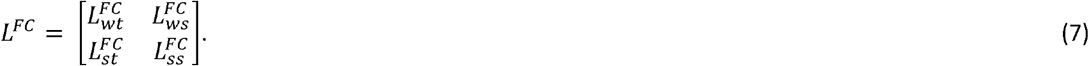

In *M^FC^* the relative contribution of the updated experimental associations to the pre-existing associations is controlled by the parameter *γ^FC^*:

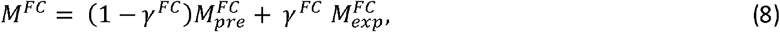

whereas the pre-experimental and experimental components of *M^CF^* do not have this trade-off:

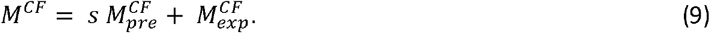

Given that the semantic associations are stored in 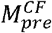 as described above, the *s* parameter thus controls the relative contribution of semantic associations to the experimentally formed (temporal and emotional) associations.

In the item layer, the emotional source units are set solely based on whether the currently presented item is emotional or neutral. In the context layer, the emotional source units are a recency-weighted sum of past emotional states, such that if emotional items were presented in the more recent past, then the emotional context unit will have a greater strength.

After all list items are presented, if there is an end-of-list distractor, it is simulated in the model by again updating c according to Equation 2. In this case, the distractor is a single item and updates temporal context with value *β_dist_*. Like the disruption to temporal context between types of emotional context, this disruption item is not incorporated into the association matrices and cannot be recalled.

At the time of recall, cuing with the context associated with the final list item *i*, **c**_*i*_ retrieves a vector 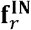. To determine which item the model recalls, 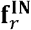 serves as the input to a leaky, competitive accumulation process (Usher & McClelland, 2001) whose value at time step *t* is determined by

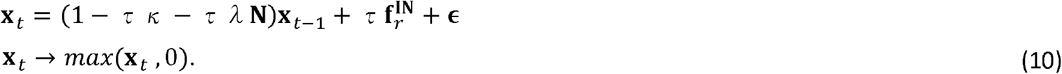

Each element of **x**_*t*_ corresponds to an element in 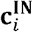. τ is a time constant and κ is a leak parameter decreasing each item by its own strength, λ is a parameter that controls lateral inhibition, by scaling the strength of an inhibitory matrix **N** which connects each accumulator to all of the others except itself, **ϵ** represents randomly distributed noise with mean 0 and standard deviation with model parameter η.

The process in Equation 7 runs iteratively until one of the accumulating elements crosses a threshold or until the recall period is over, determined by a fixed number of time steps in *t*. This equation is updated until one of the accumulating elements surpasses a threshold (here, set to 1), or until the number of time steps exceeds the amount of recall time that the model is allotted for the recall period. If an element surpasses the threshold, its corresponding item is recalled. The item is re-presented to the model, updating context according to Equation 2. Whereas the drift rate for temporal context varies between encoding and recall (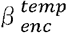 and 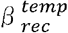 respectively) for simplicity the drift rate of the emotional context region is held constant between encoding and recall (i.e. 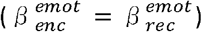. Once context is updated, this activates a different set of features on 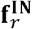, and time permitting, the recall process in Equation 10 begins again.

Below we discuss two variants of eCMR, the category-only variant, and the attention-category variant. In the latter, one additional parameter (ϕ_*emot*_) was used to model the effect of emotion on attention through an increase in the strength of association between the item features and the source context (*M^CF^*). The association strength was increased via ϕ_*emot*_ such that experimental context-feature associations were updated:

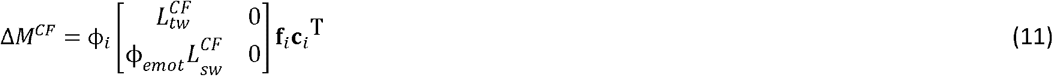

Note that Equation 11 are identical to Equation 4 if ϕ_emot_ = 1, and requires ϕ_*emot*_ > 1 in order for emotional items to benefit from stronger associations and thus improved recall.

### 2.3 Simulations of the emotional list composition effect

#### 2.3.1 Simulation 1 - the category-only variant of eCMR

We first examined simulations of the *category-only variant* of eCMR, which treats emotionality as a category such that neutral items belong to a single category and emotional items belong to another. Because emotional items are not treated specially, we did not expect to be able to reproduce the emotional memory advantage here. This variant shows how emotion is integrated into the structure of CMR, and provides a comparison that reveals the effect of attention in the next variant.

This variant of eCMR assumes, following CMR, that a change in a non-temporal context sub-region (here, a change between neutral and negative emotional contexts) disrupts the temporal context. In this way, temporal context is updated and all items studied prior to the novel item become less accessible. Thus, when two successive items are not from the same emotion category, temporal context is updated according to Equation 2. This “disruption item” only updates the temporal context sub-region, is not incorporated into the association matrices, and unlike list items, cannot be recalled. Given that a change in emotional category is qualitatively different than a presented list item, it updates temporal context according to a different model parameter, *d*, rather than 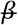 (Polyn et al., 2009). Whereas in Polyn and colleagues’ study the change in source context was a change in the externally-imposed orienting task, here the change in source context reflected a change in emotional category of the item, and concomitant internal operations. Accordingly, we set the value of the disruption parameter *d* to a lower value than that used by Polyn and colleagues (TABLE 1). The strength of semantic associations among items from the same category was simulated as **20%** stronger than the semantic associations across categories. This remained constant across all of the simulations in all sections. As mentioned in section 2.2, temporal organisation effects in the empirical data were not pronounced. In order to align model predictions of temporal contiguity with those of the empirical data, additional parameter values were altered compared to those used by Polyn and colleagues: we decreased the value of 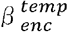, and decreased the weight the model assigns to temporal, compared to semantic, associations by decreasing 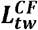.

This value of 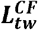 was fixed throughout this section and in each of the other sections, but varies across sections, because each of them concerns different empirical results, obtained with lists that varied in their cohesiveness. Other than that most model parameters were fixed across all of the simulations we report (Tables 1–2); we only vary parameter values to expose the machinery of the model, and this is discussed explicitly.

##### Error! Reference source not found

shows that the *category-only* variant predicted decreased memory in mixed, compared to pure lists. This decrease was due to the frequent disruptions to the temporal context during the encoding of mixed lists, which decreased the strength of associations between items, and the degree to which items were able to promote each other’s recall. In contrast to the empirical results, though, the shared emotional context of emotional items did not benefit their recall over that of neutral items. The failure of this variant to account for the list-composition effect was expected, as explained above, and led us to include emotional biases in attentional processing in the next simulation.

#### 2.3.2 Simulation 2 - the attention-category variant of eCMR

Simulation 2 was identical to Simulation 1 apart from the value of parameter ϕ_*emot*_, which was increased in order to model the emotional modulation of attention (section 2.2, equation 11).

This choice was motivated by the way attention was implemented in CMR. CMR assumes that primacy items draw attention to both experimental contexts of the presented items, i.e., temporal and source contexts. The additional attention to the earliest items in the list is implemented by increasing the strength of association between item features to temporal and source contexts (equation 5). Although emotional items might also benefit from enhanced attention in the same way, they might only draw attention to their distinctive emotional (source) context, and not their associated temporal contexts. We can distinguish between these possibilities here by asking whether emotional items modulate item features to temporal context and/or item features to source context. Because increased association to the temporal context should be manifested as increased temporal clustering scores in recalling emotional, compared to neutral lists, and there was no evidence for this in the empirical data (section 2.1), increased attention to emotional items was implemented here as increased associations between emotional items and their emotional source context.

This variant of the model simulates an equivalent increase in attention to emotional and neutral stimuli in both mixed and pure lists. Because emotionality was modelled with a binary code all emotional items in pure lists were modelled as equally attended and none should draw more attention than another. In reality, of course, some items will be more emotional to participants than others, attract more attention, and would be more likely to win the competition for retrieval during free recall.

##### Error! Reference source not found

shows that, crucially, Simulation 2 yielded an emotional list-composition effect. The comparison between simulations 1 and 2 demonstrates that preferential attention to emotional stimuli, which increases contextual binding, is necessary to the model’s ability to simulate this effect. The simulation predicted a recall advantage for emotional items over neutral ones in mixed, but not pure lists, despite equivalent attention advantage for emotional items in both types of lists. This result clarifies that differential attention alone is not sufficient to explain the decrease in memory for neutral stimuli in mixed lists. Instead, encoding and retrieval mechanisms interplay here such that the attention-dependent advantage of emotional items is more evident under conditions of competitive retrieval (in mixed lists). Section 2.4 unpacks the dynamics of retrieval to show how competition at retrieval works to favour emotional items in mixed lists.

### 2.4 The contribution of retrieval competition to the emotional list-composition effect

In this section, we unpack the retrieval mechanism of the *attention-category* variant of eCMR. We begin by showing that this model, but not the *category-only variant*, predicts an earlier output of emotional compared to neutral items throughout the recall of mixed lists, because of their stronger connection to the source context of the test (2.4.1). This pattern agrees with the empirical data. Simulation 2R shows that this early advantage is due to differential encoding of emotional items, even without further contributions from retrieval mechanisms (2.4.2). Simulations 2O, 2N and 2E explore the role of test context bias (2.4.3). They show that such bias does contribute to the advantage of emotional items, especially early on in the recall period, but that emotional items are recalled more frequently even when this bias is eliminated. This simulation therefore serves to reveal the contribution of retrieval dynamics to emotionally-enhanced memory in the model.

Early advantage influences retrieval by producing output interference. It changes the context of recall to be more emotional, and thus promotes recall of additional emotional items, and decreases the similarity between the temporal contexts of encoding and retrieval for neutral items. We present results from a new experiment to show that output interference cannot, on its own, account for enhanced recall of emotional items in mixed lists (2.4.4). To reveal how the competition between emotional and less-competitive (neutral) items during recall gives rise to enhanced emotional memory we analyse transitions during the recall, comparing predicted and empirical transitions (2.4.5). This analysis demonstrates that several dimensions of similarity multiply determine the emotion memory advantage in mixed lists.

#### 2.4.1 Output order effects during recall of mixed lists

In eCMR emotional stimuli are more strongly connected to the encoding context, which they all also share. This should give them an advantage at the first recalled position and throughout the early portion of the recall period. When an emotional item is recalled it should promote the recall of items similar to it - a self-perpetuating effect. Because emotional items share their emotional context they should promote each other’s recall. For these reasons we should expect that emotional items will be recalled earlier than neutral ones. We examine the predictions from the two variants of eCMR for recall output and found that only the *attention-category* variant predicted an earlier output of emotional items in mixed lists (Figure 4). The analysis of empirical output order effects, depicted in Figure 4 and those of Siddiqui and Unsworth (2011), showed that emotional items are recalled earlier than neutral ones, confirming the prediction of eCMR. In the following sections, we explore other aspects of the retrieval that add to the memory advantage of emotional items in mixed lists.

**FIGURE 4.**
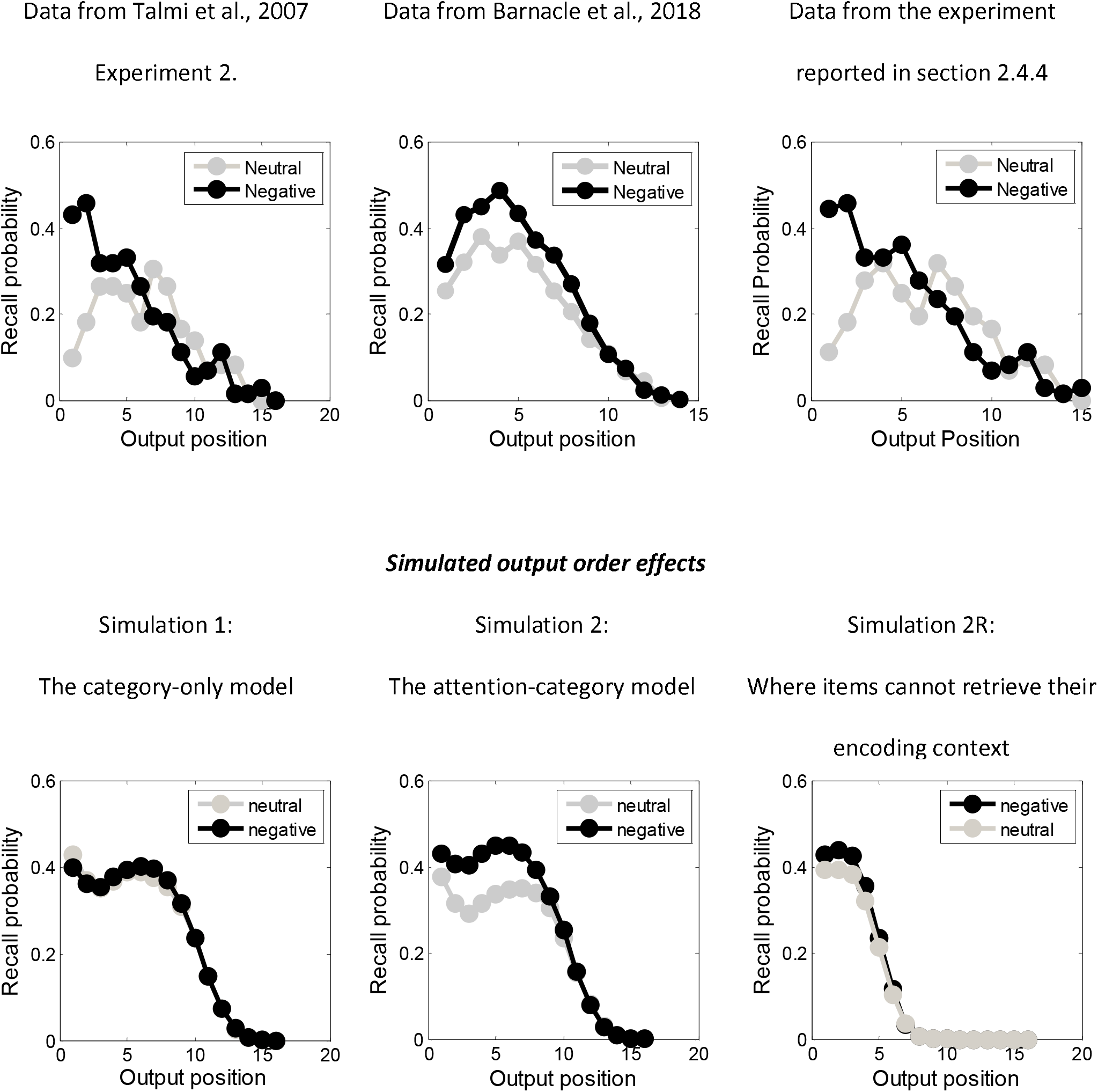
Earlier output of emotional items than neutral items during the recall of mixed lists. **Top.** Empirical output order effect. The probability of recalling items from each emotional category is plotted as a function of recall position, starting with the very first item recalled. If all of the items recalled in the first recall position were emotional, the free recall probability of emotional items would be 1 and the probability of recalling neutral items would be 0. In practice, the total probability of first recall (across both emotional and neutral items) is less than 1 because recalling non-list items, buffer items ((the first two items in the Talmi et al. and Barnacle et al.’s experiments) and other list items (random-neutral items in the Talmi et al. experiment) is scored as 0. **Bottom.** Simulated output order effects.

#### 2.4.2 Simulation 2R: Recall of mixed lists without retrieving the encoding context

Because emotional items are more strongly associated with the temporal context of the test, they should already have an advantage when recall commences, in the first output position. This advantage will influence the remainder of the recall period, even without any further influence of other aspects of the retrieval machinery of eCMR. To see this, Simulation 2R used all the same parameter values as Simulation 2, but eliminated the ability of recalled items to retrieve their study context by setting the parameters that update context during recall, 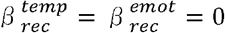. This simulation reveals, therefore, how stronger binding to the source context translates to early recall advantage in the simulation. Note that overall recall in Simulation 2R was expected to be poorer, because the ability of each recalled item to retrieve its encoding context is a core aspect of retrieved context models, so eliminating it was expected to impair memory substantially.

##### Error! Reference source not found

plots the probability of recall in Simulation 2R as a function of output order (at the bottom rightmost panel), together with the results from Simulations 1 (bottom leftmost) and 2 (bottom middle), which were discussed above. The comparison between all three simulations shows that in all of them the model recalls approximately 40% of the items. As expected, the very first recall in simulation 2R was not affected by the model’s ability to retrieve the encoding context, because this recall only depends on the initial test context. The plot also shows that in that simulation emotional items have an advantage in the first recall position, and that even without any further contributions from retrieved encoding context, this advantage extended to output positions 2 and 3. This dovetails with the predicted average recall data, depicted in **Error! Reference source not found.,** where a small emotional list composition effect can clearly be detected. In summary, this simulation reveals that preferential encoding produces an advantage for emotional items in mixed lists; comparing simulations 2 and 2R reveals how retrieval processes may enhance this advantage.

#### 2.4.3 The contribution of retrieval dynamics to emotionally-enhanced memory

The early context of the test is influenced by the last items participants saw. If the source context is more emotional than it is neutral, it would favour retrieval of emotional items, and vice versa. But because emotional items are more strongly bound to the context, an emotional context bias during the test will exert a greater influence on recall than a neutral bias. Here we explore how context bias influences the advantage of emotional items in mixed lists.

We simulated an emotional or a neutral retrieval ‘state’ by including a distractor item at the beginning of the recall period. Distractor items update the temporal context but not the association matrices, and thus are not, themselves, candidates for recall (Sederberg et al., 2008; Siegel & Kahana, 2014). We interpolated distractor item with a null (Simulation 2O), emotional (Simulation 2E), or neutral (Simulation 2N) context just before the recall period. The magnitude of influence of this distractor item was controlled by *β_emot_*.

In Simulation 2O we can see a small advantage for emotional items. In Simulation 2E biasing the retrieval context with an emotional distractor – essentially, rendering it more emotional - increased the likelihood of recalling emotional items. Biasing the test context away from emotionality with a neutral distractor, in Simulation 2N, reversed the bias, resulting in increased first recall of neutral items. Crucially, because each recalled item retrieves its own source context, which dilutes the impact of the bias, no bias was observed in the second output position, and the emotional bias reappeared in the third output position (Figure 5).

#### 2.4.4 The emotional advantage in mixed lists does not depend solely on interference at retrieval

The earlier recall of emotional items could create output interference, which could produce the emotional memory advantage in mixed lists even without the other mechanisms postulated in eCMR. To examine the impact of output order on the emotional memory advantage in mixed lists we conducted a new experiment, described in **APPENDIX 1APPENDIX 2.** Retrieval of mixed lists of negatively-valenced emotional and neutral pictures was manipulated by asking participants to recall emotional or neutral items first (the ‘emotional first’ and ‘neutral first’ conditions). The crucial result for this section concerns the data from the ‘emotional first’ group, who were instructed to recall the emotional pictures first, and then the neutral pictures; and the ‘neutral first’ group, who were instructed to recall the neutral pictures first, and then the emotional pictures. Let us examine the predictions of eCMR for these conditions. An emotional memory advantage should still be obtained even when neutral items are recalled first, because emotional items compete for recall with items that are less strongly bound to the encoding context. While output interference is expected to decrease memory for both emotional and neutral stimuli, eCMR predicts that an emotional advantage should be present both when we compare the first recalls of each group (the recall of emotional items in the ‘emotional first’ group and the recall of neutral items in the ‘neutral first’ group), and when we compare the second recall of the two groups (the recall of neutral items in the ‘emotional first’ group and the recall of emotional items in the ‘neutral first’ group). By contrast, a greater difference in emotional versus emotional stimuli when comparing the first recall test to the second will support the hypothesis that this advantage depends on output interference. If the emotional memory advantage depends solely on output interference, then the emotional advantage should be greatly weakened in the first recall of the ‘emotional first’ and ‘neutral first’ groups, where such suppression is prevented, compared to the second recall of those groups, which more closely mimics the ‘natural’ uninstructed condition. The logic of eCMR dictates, however, that because emotional items are more strongly associated to the encoding context and to each other, the emotional advantage in mixed lists would remain even when output interference is reduced or eliminated – as long as emotional items compete for recall with non-emotional items. In this section, we describe empirical evidence in support of this hypothesis.

We observed a significant emotional memory advantage of equivalent magnitude in the first and the second recall (*FIGURE 6. Figure 5*). The analysis of the instructed conditions used two independent sample t-tests to compare recall of emotional and neutral items across the ‘emotion first’ and ‘neutral first’ groups. The magnitude of the effect of emotion on memory was equivalent and large in the first recall test (comparing recall of emotional items in the ‘emotion first’ group to neutral items in the ‘neutral first’ group), t(34)=2.34, p<.05, Cohen’s d=0.80, and in the second recall test, (comparing recall of neutral items in the ‘emotion first’ group to recall of emotional items in the ‘neutral first’ group), t(34)=2.20, p<.05, Cohen’s d=0.76. These results agree with the prediction of eCMR and going against the interpretation of the data as due solely to output interference.

**Figure 5.**
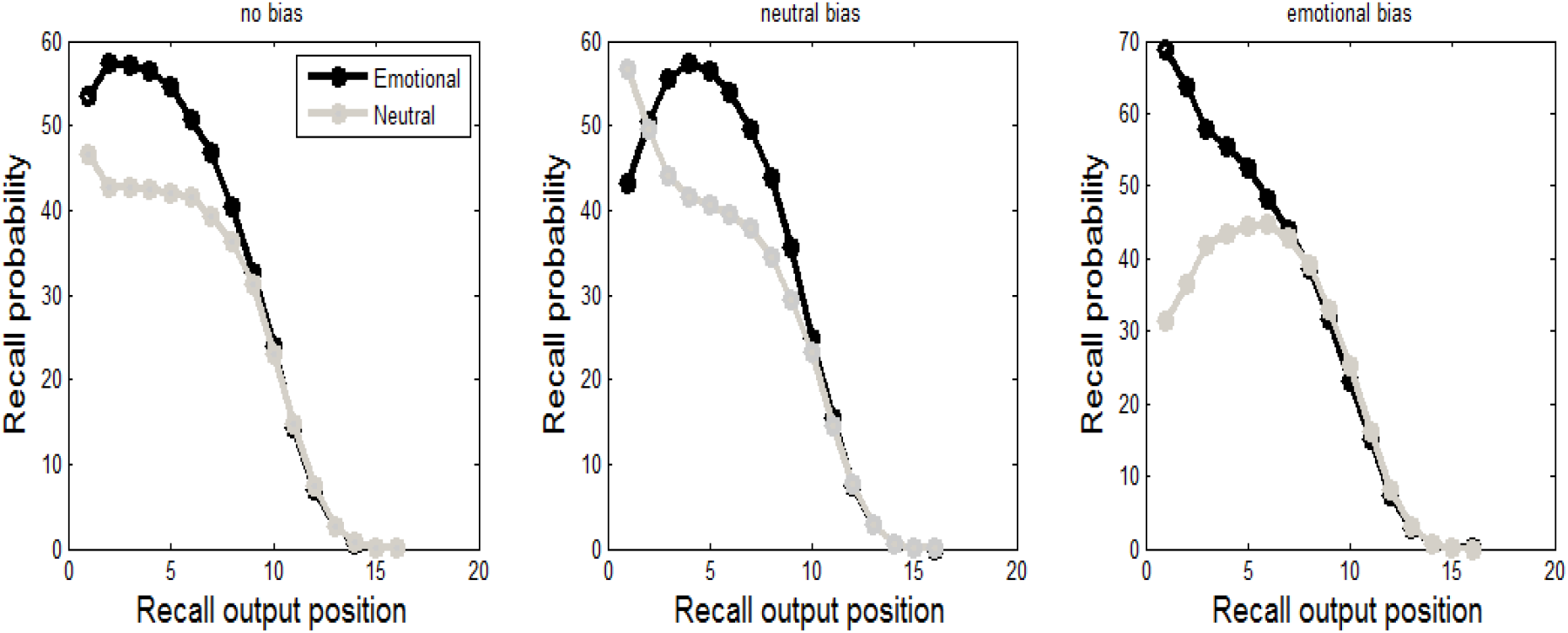
Free recall of emotional and neutral items, encoded in mixed lists, as a function of the emotional context of the test. **Left.** No bias (Simulation 20). **Middle.** Neutral bias (Simulation 2N). **Right.** Emotional bias (Simulation 2E).

**FIGURE 6.**
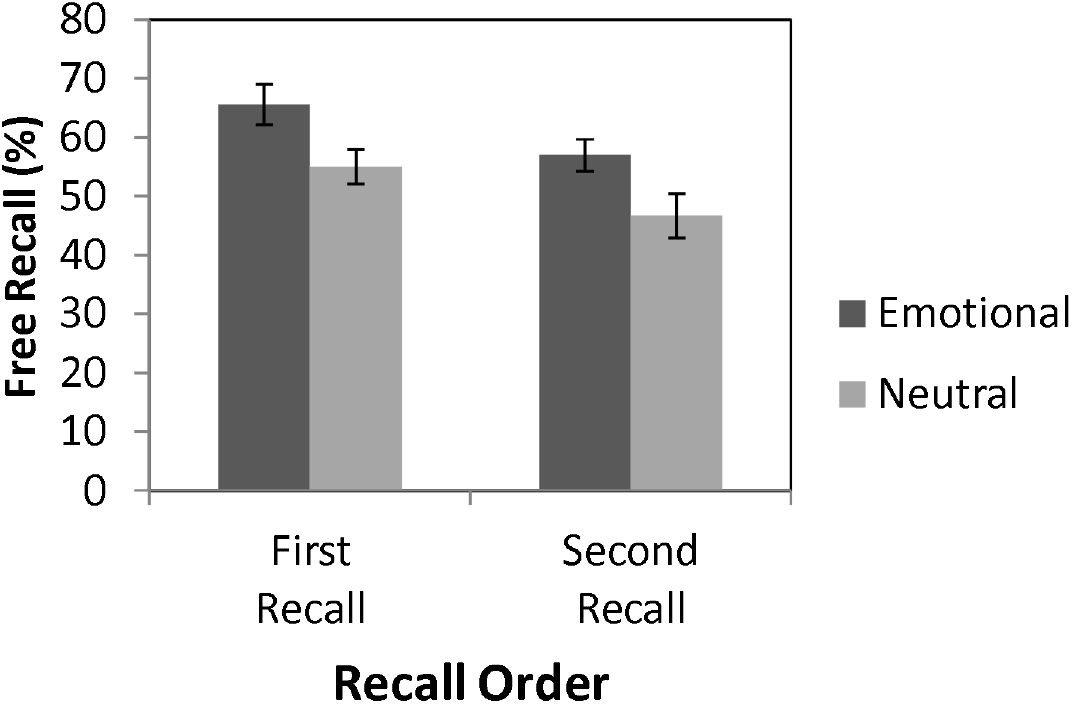
Empirical data from the instructed recall experiment (Appendix 2) Free recall of the ‘emotion first’ and neutral first’ groups is plotted as a function of the test period (the first or second recall). The two leftmost bars plot the recall from the first test period in each of these two groups (the emotional list in the ‘emotion first’ condition and the neutral list in the ‘neutral first’ condition). The two rightmost bars plot recall from the second test period in each of these groups (the neutral list in the ‘emotion first’ condition and the emotional list in the ‘neutral first’ condition).

#### 2.4.5 Recall transitions in the attention-category variant of eCMR

To fully understand the interaction between attention to emotion at encoding and the retrieval machinery of eCMR we analysed the types of transitions the model made between successive recalls in Simulation 2, following the same methodology used by Polyn and colleagues (2009, 2011), as described in section 2.1. The descriptive results are provided in TABLE 4.

Temporal clustering within category was greater than chance in the empirical data and in the model, reflecting both the shared emotional features, as well as the close temporal proximity, of these items. Between-category temporal clustering is an index of the contribution of temporal associations between items even when they do not share the same emotional context. The model over-predicted between-category temporal clustering; it was as high as within-category temporal clustering in the simulation, but at chance in the empirical data. This discrepancy presents a limitation of the model which could be examined in future work. Next, we examined the emotional clustering scores. eCMR naturally predicts emotional clustering effect because recall of an item leads to retrieval of its associated context states, including its emotional context. As in the empirical data, transitions based on shared emotional context in the model were more likely than what is expected at chance. We also saw a relationship, such that across simulated “subjects”, recall probability of emotional items was greater when the emotional clustering score was high, while recall of neutral items correlated negatively with emotional clustering. This relationship was not observed in the empirical data, but as stated above, these data had limited power, and it is difficult therefore to interpret these null effects. Lastly, we considered how transitions between successive recalls were influenced by semantic organisation. eCMR, like CMR, predicts that like temporal and emotional dimensions of similarity, semantic similarity should also contribute to recall. A semantic clustering score can be defined in a way analogous to the temporal clustering score, except rather than defining any pair’s similarity based on their absolute temporal lag, similarity was defined based on the estimated semantic association in the simulation. The semantic clustering score in the model was greater than chance. We could not examine this in the empirical data because of the absence of an appropriate complete matrix of semantic similarity scores, but the model’s prediction can be examined in future research.

Taken together, these results show that in both the empirical data (section 2.1) and in Simulation 2, the dynamics of recall of mixed lists depend jointly on all of the associations that encoded items share. When an emotional item is recalled, it would most likely promote the recall of another emotional item. This is also true, of course, for neutral items; but because emotional items are recalled first, and promote recall of other emotional items via their shared emotional context that they are more strongly connected to, their recall benefits more from the propensity to cluster around the emotional dimension.

### 2.5 Comparison of eCMR and the item-order account

eCMR formally embodies some of the same rationale of the item-order account, which was developed to explain non-emotional list composition effects for atypical compared to standard items (McDaniel & Bugg, 2008). Like retrieved context models, the item-order account recognizes the central role of items’ temporal order. Both the item-order account and eCMR assume that recall performance reflects a trade-off between the contributions of temporal context and other item characteristics. But the temporal order in the item-order account plays a different role than the temporal context in eCMR.

In the item-order account, memory for the order of list items helps recall only when it is attended during encoding; and attention to order decreases when it is otherwise engaged, for example when more attention is allocated to processing the items’ identity (McDaniel and Bugg, 2008). When lists include unusual items participants elaborate on their identity and pay less attention to order. Therefore, attention to the temporal order of atypical items in pure lists is limited; it is greater when lists include some standard items (mixed lists); and greatest in pure lists of standard items. While attention to the identity of atypical items maintains their good recall regardless of list composition or order memory, any extra attention to the identity of atypical items takes away from the attention participants allocate to the order of standard items, resulting in their decreased recall.

In the item-order account attention to the temporal order helps recall by supporting forward recall transitions. Therefore, to support the item-order account, McDaniel and Bugg (2008) examined the “input-output correspondence index” (Asch & Ebenholtz, 1962), which is defined as the proportion of transitions made between successive items at lag=+1 (thus, the input and output order correspond to one another). In agreement with the predictions of the item-order account this index was lowest when participants encoded pure unusual lists, highest when participants encoded pure standard lists, and at a middling level when they encoded mixed lists (e.g. McDaniel, DeLosh, & Merritt, 2000). We computed the same index for the empirical datasets we simulated in this section, and found that the same was true for Talmi et al. (2007: pure emotional lists: M=.07, SD=.14; mixed lists: M=.10, SD=.09; pure neutral lists: M=0.14, SD=.13). In the Barnacle et al. dataset the input-output correspondence index was the same for pure emotional lists (M=.11, SD=.07) and mixed lists (M=.11, SD=.10), but, as predicted by the item-order account, higher for pure neutral lists (M=0.17, SD=.12). Also in agreement with the item-order account, the difference between the input-output correspondence indices in the recall of pure lists was significant in both datasets. To make the links between eCMR and the item-order account explicit we computed the input-output correspondence indices for simulated data in Simulation 2, but did not observe much difference in the input-output correspondence index between list types (pure emotional lists: .15, mixed lists: .16, pure neutral lists: .16). The item-order account therefore predicts an aspect of the empirical data that eCMR does not. The parameters we selected for the simulations we reported in this section were based on the best-fit set in Polyn et al. (2009), so it is possible that a different selection of model parameters would allow eCMR to fit this result, but additional research with more powered empirical data is required to decide exactly what recall dynamics look like for the emotional list composition paradigm. eCMR did capture the above-chance temporal clustering scores in the empirical datasets, which collapse across all transitions, including the lag=+1 that the item-order account focuses on.

When we come to evaluate the two models in terms of their account of the list composition effect, perhaps the most important consideration is that the attention-category variant of eCMR can reproduce the list-composition effect without making the assumption that the item-order account makes - that attention to neutral items during the encoding of mixed lists is decreased, compared to the attention they receive in pure lists, and which contradicts the empirical data (Barnacle et al., 2016; Barnacle & Talmi, 2016; Talmi & McGarry, 2012). Unlike the item-order account, eCMR does not mandate a particular weighting of context dimensions. While reliance on the source context could decrease temporal contiguity effects (Polyn et al., 2011, 2009), in the direction predicted by the item-order account, this did not happen in the empirical studies we reported here (TABLE 4). Overall, eCMR is a more comprehensive model, and can account for more aspects of the data than the item-order account; for example, only eCMR captured recall transitions based on source context, which were observed in the empirical data.

### 2.6 Summary

The empirical and simulation results in this section deconstruct the emotional memory advantage to its constituting elements. In both of the eCMR variants explored in section 2 emotional arousal was modelled as an emotional feature of items that updates an emotional context sub-region. An emotional (relative to a neutral) source context did not confer particular advantages. In the *category* eCMR variant memory for both emotional and neutral items in mixed lists suffered from the disruption to the temporal context through the frequent alteration between item types. In the *attention-category* eCMR variant emotional items were modelled as systematically attended more, thus bound more strongly to their source (emotional) context. This improved their chances of winning the competition for retrieval. This variant captured the pattern of average free recall and the dynamics of the emotional list-composition effect.

The success of eCMR to capture the emotional list composition effect depended on a number of retrieval mechanisms that interplayed with the additional attention towards emotional items during encoding. Overall recall success in eCMR depended on the similarity between the retrieval context and the temporal, semantic, and (emotional) source context of encoded items. Emotional items in mixed lists were more likely to be recalled first because they were more strongly connected to the encoding context, and thus won the competition for recall. Once one emotional item was recalled, the shared emotional context between these items rendered additional emotional items more accessible. We note that increased cohesiveness among emotional items, when this is not controlled experimentally, would also contribute in the same way, but here we equated the similarity of emotional and neutral items, in keeping with the empirical data we simulated. Because emotional items tend to be recalled early and promote each other’s recall they act as a source of interference, making it more difficult for participants to retrieve neutral items. We showed that empirically, an emotional advantage is observed even when output interference is eliminated. The simulations clarified that an emotional advantage appears even when the initial source context of the test, and therefore the probability of first recall, was biased towards recall of neutral items.

In pure lists enhanced attention to emotional stimuli still increases their binding to emotional source context, but because the increased level of binding is equivalent for all of the emotional items, the competition for retrieval is affected by emotionality less than in mixed lists. Therefore, although encoding processes could be very similar in pure and mixed lists, the advantage of emotional items in free recall would be expressed more strongly in mixed lists. Taken together, there are several processes in place that give rise to the emotional advantage in free recall of mixed lists, rendering the emotional list-composition effect a robust phenomenon, well-described by eCMR.

## 3. eCMR can account for increased emotion-enhanced memory after a delay

A paradigmatic finding in the emotional memory literature is that of a steeper forgetting of neutral compared to emotional stimuli (Yonelinas & Ritchey, 2015). In reports where the memory advantage is greater in a delayed test the advantage may be smaller immediately and grow with delay (e.g. LaBar & Phelps, 1998); there may be no immediate advantage (e.g. Sharot & Yonelinas, 2008; Sharot & Phelps, 2004); or, in a few, classic, reports, emotional stimuli may be remembered less well than neutral stimuli immediately, but better later (e.g. Butter, Training, & Kaplan, 1970; Kleinsmith & Kaplan, 1964). These studies used many different methodologies to trigger arousal and to measure memory (free recall, cued recall and recognition), which could well account for some of the discrepancies. A systematic review of this issue is certainly warranted, not only to explain the discrepant immediate effects, but also to provide a more solid basis for the consensus in the emotional memory literature that emotion-enhanced memory effects are greater after a delay. Because the immediate effects of emotion are discrepant they are often disregarded, as are, by extension, interpretations of delayed effects that refer to encoding or retrieval mechanisms. Instead, the emphasis in the emotional memory literature has been the attenuated forgetting of emotional stimuli, and how this effect is implemented through neuromodulation (Dunsmoor, Murty, Davachi, & Phelps, 2015; Patil, Murty, Dunsmoor, Phelps, & Davachi, 2017).

Readers familiar with work on the neurobiology of memory consolidation may conclude, therefore, that eCMR has no hope of accommodating attenuated forgetting of emotional items, because it lacks any mechanism to mimic consolidation effects. Yet although retrieved context models have been used to account for memory effects on time scales more commonly used in human memory experiments, such as a single session, some model variants have been developed to explain consolidation-related effects. Those operationalise the retention interval through increased interference, or through degraded context reinstatement (Howard, Kahana, & Wingfield, 2006; Sederberg, Gershman, Polyn, & Norman, 2011; Sederberg et al., 2008). In this section, we use eCMR to simulate a prolonged retention interval, and examine how delay might alter the emotional advantage. Our purpose was to illuminate which cognitive processes influence changes in the magnitude of the emotional memory advantage over time. eCMR is agnostic as to how such changes may be realised at the level of neurobiology.

Deciding on the most appropriate way to simulate memory delay is beyond the scope of this paper. Further research is needed to decide on the value of each of the parameters that might change with delay in a particular set-up. In order to examine the predictions of eCMR for delayed effects of emotion on memory here we utilised a combination of the approaches implemented in retrieved context models elsewhere, as described below. Therefore, Simulation 3 could be seen as a proof of concept for greater effects of emotion in delayed tests, rather than a commitment for a particular implementation. To show this most clearly, Simulation 3 implements a classic experiment where emotional items had an increased free-recall advantage in a delayed test, compared to an immediate test (LaBar & Phelps, 1998).

### 3.1 A description of empirical data from LaBar and Phelps, 1998

LaBar and Phelps (1998) studied healthy controls as well as patients with unilateral temporal lobectomy. Participants incidentally encoded a single mixed list of 20 low-frequency neutral words and 20 taboo words. Four buffer words were also included, two in the beginning and two at the end of the list. During encoding participants rated how arousing the words were. A free recall test was administered immediately and after a 1-hour filled delay. The researchers found an immediate increase in the number of taboo words recalled, which they attributed to the increased inter-relatedness of these words compared to the neutral words. Strikingly, participants forgot fewer taboo words, so that their memory advantage was greater after one hour than it was immediately, an effect they attributed to the effect of arousal on consolidation (*FIGURE 7*). The impact of this work, which has been cited over 200 times (retrieved from Web of Science, July 2018) is due, in part, to the finding that the participants with temporal lobectomy forgot taboo and neutral words at the same rate, in agreement with the hypothesis that the attenuated forgetting of emotionally arousing material is driven by the amygdala (LaBar & Cabeza, 2006).

**FIGURE 7.**
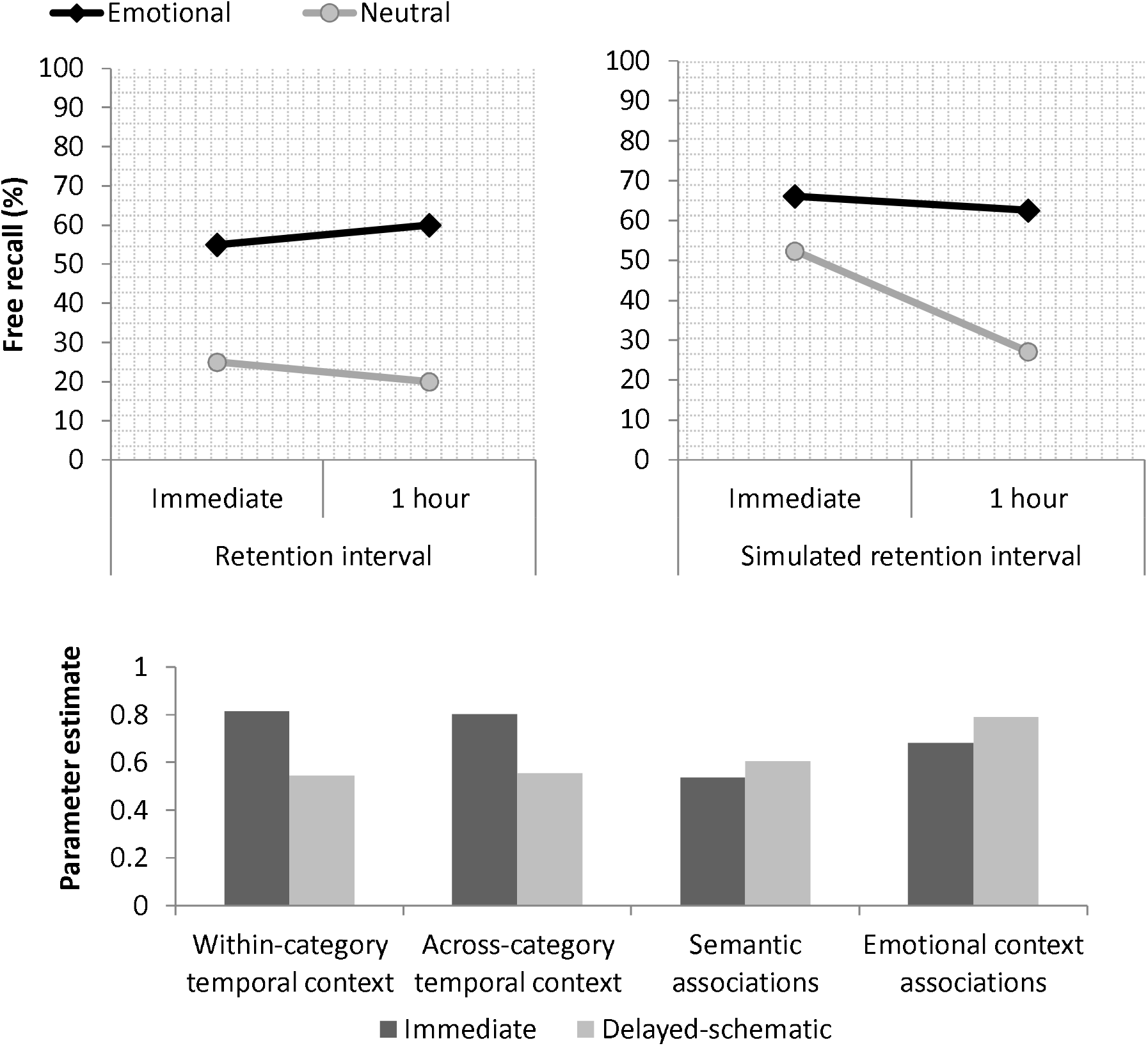
Forgetting effects. **Top:** Emotional memory advantage as a function of time delay. **Left:** Original data adapted from Figure 2 in LaBar and Phelps (1998). Black: taboo words. Grey: neutral words. **Right.** Predictions of Simulations 3I and 3D. **Bottom.** Parameter estimates for predicted clustering effects in Simulation 3l and 3D.

Notably, although the results suggested an *increase* in emotional memory after a delay, in the empirical data the interaction between item type and time was significant, but not the simple effects. While steeper proportional forgetting of neutral items in free recall has been reported by others (e.g. Bradley, Greenwald, Petry, & Lang, 1992; Christianson & Loftus, 1987; Christianson, 1984; Farley, 1973), there are also contradictory reports (e.g. Maltzman, Kantor, & Langdon, 1966; Schwabe, Bohringer, Chatterjee, & Schachinger, 2008). In particular, a meta-analysis of this issue could establish not only the magnitude of emotion-modulated forgetting and the conditions that influence it, but also address potential file-drawer biases due to the strong theoretical predictions of emotion-modulated forgetting. Therefore, while Simulation 3 reveals the conditions that allow such an effect to become manifest, it is not known at present how typical these conditions are in reality.

### 3.2 Simulating prolonged retention intervals in eCMR

One way of simulating a retention interval is to simulate a ‘distractor’ item by inserting a temporal context disruption at the end of the encoding phase (Siegel & Kahana, 2014; Sederberg et al., 2008). While this approach has been successfully utilised to capture the effect of time delays in the order of seconds to minutes, it may not quite capture longer delays, such as the hours-to-weeks delays employed in the emotional memory literature. An additional problem with this approach is that in eCMR, once one item is recalled, this leads to retrieval of the item’s encoding contexts and effectively allow the model to ‘jump back in time’, so that the effects of the retention interval would be limited to the earliest stages of retrieval. Therefore, in addition to simulating post-list distraction, which does probably occur, we modelled the retention interval as decreasing the ability of recalled items to retrieve the temporal context of their encoding. We decreased the overall retrievability of the temporal context by decreasing 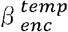 an approach which has previously been employed successfully to capture the effects of aging on the recency and contiguity effects (Howard et al., 2006). In addition, we assumed that, after a delay, each item is less well associated with its own temporal context, so that it cannot retrieve it very well. For this purpose we decreased 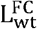 and 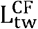, an approach taken by Sederberg and colleagues to model part of the deficit in people with anterograde amnesia, which is characterised by poor delayed memory (Sederberg et al., 2008).

Because in eCMR all context dimensions support recall jointly, a decrease in the reliance of the model on the temporal context during retrieval by necessity increases its reliance on non-temporal dimensions, such as the semantic and the emotional contexts. Therefore, the changes we implemented to simulate a prolonged retention interval correspond well to evidence that forgetting is not passive, but involves a transformation of memory traces so that memory for more remote events is more schematic (McKenzie & Eichenbaum, 2011; Moscovitch, Cabeza, Winocur, & Nadel, 2016). For example, animals that learned the location of a hidden platform in a Morris water maze based their trajectory more on the specific location of the platform when memory was recent, but were more sensitive to the probability distribution of platform locations when the memory was remote (Richards et al., 2014). A long retention interval is also thought to allow humans to extract generalities (Ellenbogen, Hu, Payne, Titone, & Walker, 2007) and render memory more schematic (Alba & Hasher, 1983). For example, remote memory for film clips relied more on general schemas for films and social scripts, evident in increased number of errors in recalling typical vs. atypical clips after a week (Bonasia et al., n.d.), suggesting increased relative accessibility of schema-relevant details after a delay (Sekeres et al., 2016). The literature does not, at present, specify exactly what time frame gives rise to more schematic memory performance. It is also possible that aspects of the experimental set-up, such as the stimulus set that is selected or the particular instructions participants receive, influence the strategies participants apply to the difficult task of delayed recall. Yet one prediction is clear: because non-temporal dimensions of similarity promote the recall of emotional items in eCMR, their greater influence after a delay is likely to increase in the simulated emotional advantage.

### 3.3 Simulations 3l and 3D: Immediate and delayed memory

We simulated 30 participants who each studied a list of 16 items, half of which were emotional. Note that we deliberately simulated lists that were shorter than those used by LaBar and Phelps (1998), in order to keep parameter estimates as close as possible to those used in section 2, and facilitate comparison across sections. Nevertheless, there were a few differences between Simulations 2 and 3. As in the original study the first and last two items were considered buffers and not analysed; two of these were emotional, two of these neutral, and they were allocated randomly to the 1^st^, 2^nd^, 17^th^ and 18^th^ list positions for every simulated participant. To better match the empirical setup of LaBar and Phelps, the simulated immediate test in Simulation 3I followed the encoding stage without simulating a distractor at the end of the list. Based on the discussion in the original paper, we assumed that emotional items were more inter-related to each other than the neutral items. In Simulation 3 the average semantic associations strength was 0.072, apart from among emotional items, where it was 0.09. Because LaBar and Phelps used neutral items which were not semantically related, it was likely that the semantic context played a less important role in immediate recall in their study, compared to the study we simulated in section 2. We therefore used the original value for 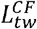, from Polyn et al. (2008).

To recap, in Simulation 3D the delayed test was modelled by (1) simulating a distractor at the end of the list (2) decreasing 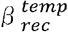, which governs the overall ability to retrieve the temporal context (3) decreasing parameters 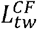 and 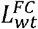, which control the associations between temporal context and temporal item features (TABLE 1). These changes weaken the associations between the item and its temporal context. Consequently, recalling one item will promote the recall of items that share its temporal associations to a lesser extent, instead promoting the recall of items that share its semantic and emotional context.

### 3.4 Comparison between Simulations 3I and 3D

Simulations 3I and 3D captured the full pattern of data described by LaBar and Phelps (1998). About half of the neutral items were forgotten after a simulated delay, while memory for emotional items persisted. Compared to the recall advantage of emotional items in Simulations 3I, in Simulation 3D the model recalled both numerically and proportionally more emotional than neutral items (*FIGURE 7*). It is well known that forgetting can be quantified in at least two different ways – the decrease in the number or the proportion of retained (Wixted, 1990); here the effects are large, and stand regardless of how forgetting is quantified. Further examination of these results suggests that increasing the value of the distractor at the end of the list, or decreasing 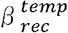 further, decreased recall of both emotional and neutral items, while the value of 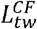 and 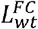 influenced the relative forgetting of these item types.

The difference between the predictions eCMR makes for immediate and delayed memory can be elucidated by analysing the predictions those simulations make for the organisation of recall (see sections 2.1 and 2.4.5 for details of the analysis method). *FIGURE 7FIGURE 9* shows that simulated delay resulted in lower temporal clustering scores, together with increased influence of emotional and semantic factors on recall. Increased reliance on the semantic context with delay is reminiscent of the empirical demonstration that semantic similarity interacts with the retention interval, such that the relatedness of the retrieval cue and the target item influenced delayed cued recall more than immediate cued recall (Schooler et al., 1997). Simulation 3D suggests that the same might be true for the emotional context.

Taken together, the simulations in this section suggest that the attenuated forgetting of emotional stimuli - or even the absence of any forgetting at all - may not be a direct consequence of the retention interval, but an indirect consequence of the effect of the retention interval on the balance between the contextual dimensions that aid retrieval of particular memories.

## 4. The emotional oddball effect

In Sections 2 and 3 we considered recall of mixed lists where the number of emotional versus neutral items was equivalent, and in section 2 also recall of pure lists that were comprised entirely of emotional items or entirely of neutral items. To fully characterize the interaction of emotion and memory in recall we should also examine what happens when lists have other ratios of emotional to neutral items. An obvious case is the emotional oddball effect, where lists are comprised solely of neutral items, with the exception of one emotional oddball. For example, in a classic paper (Ellis, Detterman, Runcie, McCarver, & Craig, 1971) participants studied lists of line drawings of everyday objects for free recall. The emotional oddball was a picture of a model from a ‘sun tanning’ magazine, presented in a middling serial position. The oddball was recalled extremely well, while memory for surrounding items suffered, compared to control lists without oddballs. This decreased memory for items presented before and after the oddball is often referred to as retrograde and anterograde amnesia, respectively. Together with excellent memory for the emotional oddball itself, this pattern is referred to as the emotional oddball effect. The emotional list composition and the emotional oddball effects resemble each other because they are observed in tasks where emotional and neutral stimuli are presented in the same list and involve neutral memory impairments, and because both are sensitive to multiple cognitive processes (Schmidt & Schmidt, 2016).

Although the similarities between the emotional oddball effect and the emotional list composition effect suggest that eCMR should be able to predict both, the emotional oddball effect nevertheless presents a significant challenge to the model. The most obvious problem is that by definition, an oddball is less similar to standard list items compared to the similarity among standard items. In retrieved context models decreased similarity would impair recall, which contradicts the robust finding, known since the experiments of von Restorff, that deviant items are recalled extremely well (reviewed in Hunt & McDaniel, 1993). There are also plenty of demonstrations that this is true for an emotional oddball (Ellis et al., 1971; Hurlemann et al., 2005, 2007; Mather & Knight, 2009; Schmidt, 2012; Strange, Hurlemann, & Dolan, 2003).

This is not the only problem. In retrieved context theory, when an item is recalled it promotes the recall of items with shared context elements, including shared temporal context. This is a core aspect of the theory, and helps it explain temporal contiguity effects in free recall (Howard & Kahana, 1999, 2002a). If eCMR could simulate the excellent recall of the oddball item, then recall the oddball will promote the recall of its nearest neighbours (oddball-1 and oddball+1). Yet this prediction is exactly the opposite of the retrograde and anterograde amnesia aspects of the empirical oddball effect, which is thought to be even more accentuated when the oddballs are emotional (Hurlemann et al., 2005, 2007; Mather & Knight, 2009; Strange et al., 2003). Admittedly, the exact pattern of oddball-induced retrograde and anterograde amnesia vary; there have been reports of retrograde without anterograde amnesia (Tulving & Thomson, 1973), anterograde without retrograde amnesia (Schmidt, 2002), or both anterograde and retrograde amnesia (Detterman & Ellis, 1972). But Schmidt and Schmidt (2016) review the literature on the emotional oddball, and conclude that amnesia effects in tasks such as immediate free recall are robust. Specifically, they suggest that anterograde amnesia effects are robust across stimulus types and types of test (recall and recognition), and that retrograde amnesia effects are observed consistently in recall tests that take place shortly after encoding, as in the cases we simulate here.

In eCMR we have a ready mechanism to implement increased attention, which could go some way to counteract the dissimilarity of the oddball to the context, and boost its recall. Yet boosting the recall of the oddball is unlikely to address the problem of anterograde and retrograde amnesia for items near the oddball, for the reasons discussed above. One possible resolution to this discrepancy between the emotional oddball effect and the generic predictions of retrieved-context theory has to do with the assumption of CMR that the temporal context is disrupted whenever item sources change, something that occurs when an emotional item is presented after a series of neutral items, or vice versa. We examine these solutions below. Our main aim in this section is to see what mechanisms permit eCMR to simulate the superior memory for an outstanding isolate, and examine which of these also impair memory for items in close serial positions. By examining what elements of eCMR need to change in order to reproduce the emotional oddball effect we may develop deeper understanding of the empirical effect.

### 4.1 Simulation 4 and 4A: Representing an oddball in eCMR

By definition, an oddball is an item that is *different* from the standard list items, and therefore likely to trigger different internal operations than the standards. This can be readily operationalised in eCMR through oddball and standards having different source contexts. Notably, so far we have only considered source contexts that are emotional and neutral, but the same approach can be used to represent other differences in the encoding of standards and oddballs, such as the difference between line drawing standards and an oddball picture, for example. Because the test context will be dominated by the source context of the standards, an oddball with a different source context should be recalled less well. Additionally, an oddball may also be less related to standard items on a certain dimension, such as its pre-experimental semantic associations. This is certainly the case for mixed lists of emotional and neutral items, where even if these two item categories are equally cohesive, they are less related to each other. Decreased semantic similarity will also hinder the recall of oddballs. Emotional oddballs would attract attention both because they are emotional and because they are unexpected; unexpected items are attended and processed preferentially (Kok, Rahnev, Jehee, Lau, & de Lange, 2012). Increased attention to the oddball will increase its link to the temporal context, help its recall, and could even counteract the impact of decreased source and semantic similarity. The balance between these effects of similarity and attention would determine how well an oddball is remembered. This balance will depend on aspects of the experimental set-up, such a type of oddball used.

We simulated lists of 15 neutral items and a single emotional item in position 9. The same list length was used in Simulation 2; this list length and the serial position of the oddball are also similar to the one used by Ellis et al. (1971). We also simulated control lists with 16 neutral items. Simulation 5 used the parameters of eCMR from section 2, apart from the degree of reliance on the semantic associations, which was mid-way between Simulation 2, where we simulated cohesive lists, and Simulation 3, where we simulated randomly-selected lists. Simulation 4 did not predict superior memory for the oddball compared to standards, but the reverse - a dip in memory for the oddball (*TABLE 5*). This Simulation is useful because it exposes the challenge that the emotional oddball effect poses for eCMR.

**TABLE 5.** *Recall (%) in Simulation 5*.

|  | *Oddball list* | *Control list* |
| --- | --- | --- |
| *Positions 1-8* | 76.22 | 75.30 |
| *Oddball (position 9)* | 29.24 | 66.10 |
| *Positions 10-16* | 63.46 | 64.66 |

In Simulation 4A we shifted the balance between the various mechanisms that implement the oddball by simulating an oddball that attracts more attention than emotional items in mixed lists. Simulation 4A captured the idealised pattern we described as the emotional oddball effect: excellent memory for the oddball itself, accompanied by worse memory for items in other list positions, compared to control lists without oddballs (*FIGURE 8)FIGURE 9*. Memory for items near the oddball was particularly impaired, a result that amounts to anterograde and retrograde amnesia effects for items near the oddball. In the next section we turn to empirical data, to reveal which of the mechanisms we used to simulate the oddball in Simulation 4A is specific to emotional oddballs.

**FIGURE 8.**
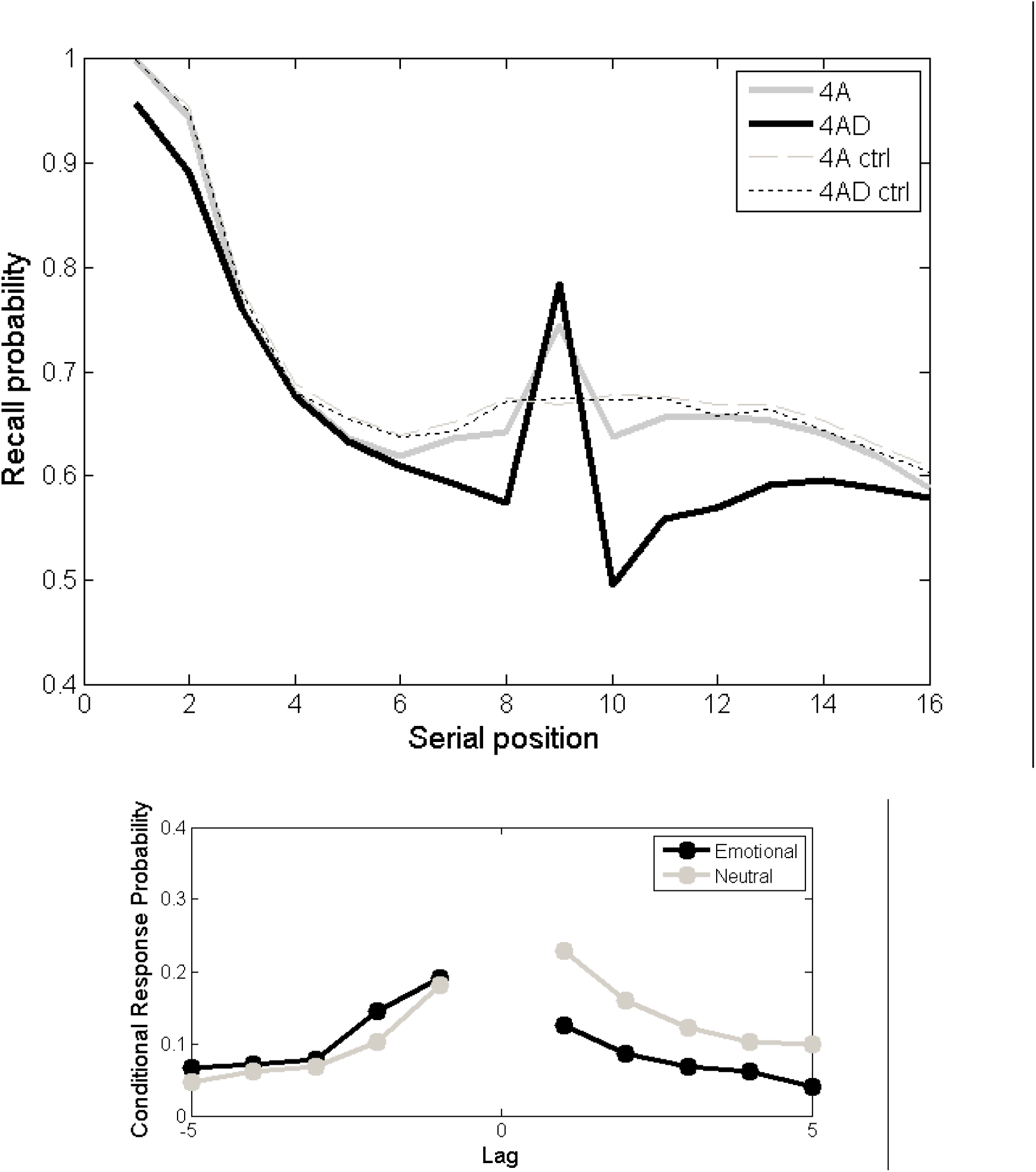
The emotional oddball effect. **Top.** Predicted recall in Simulation 4A and 4AD for lists that included a single oddball item in position 9 and for control lists comprised of standard items only, as a function of serial position. Simulations 4A and 4AD differed in the degree to which the oddball disrupted the temporal context, with 4A simulating the neutral oddball and 4AD the emotional oddball. **Bottom.** Predicted conditional recall probability as a function of lag, depicting transitions from the emotional (black, Simulation 4AD) or neutral (grey, Simulation 4A) oddball. The plot shows a decreased tendency to retrieve the item that follows the emotional oddball.

**FIGURE 9.**
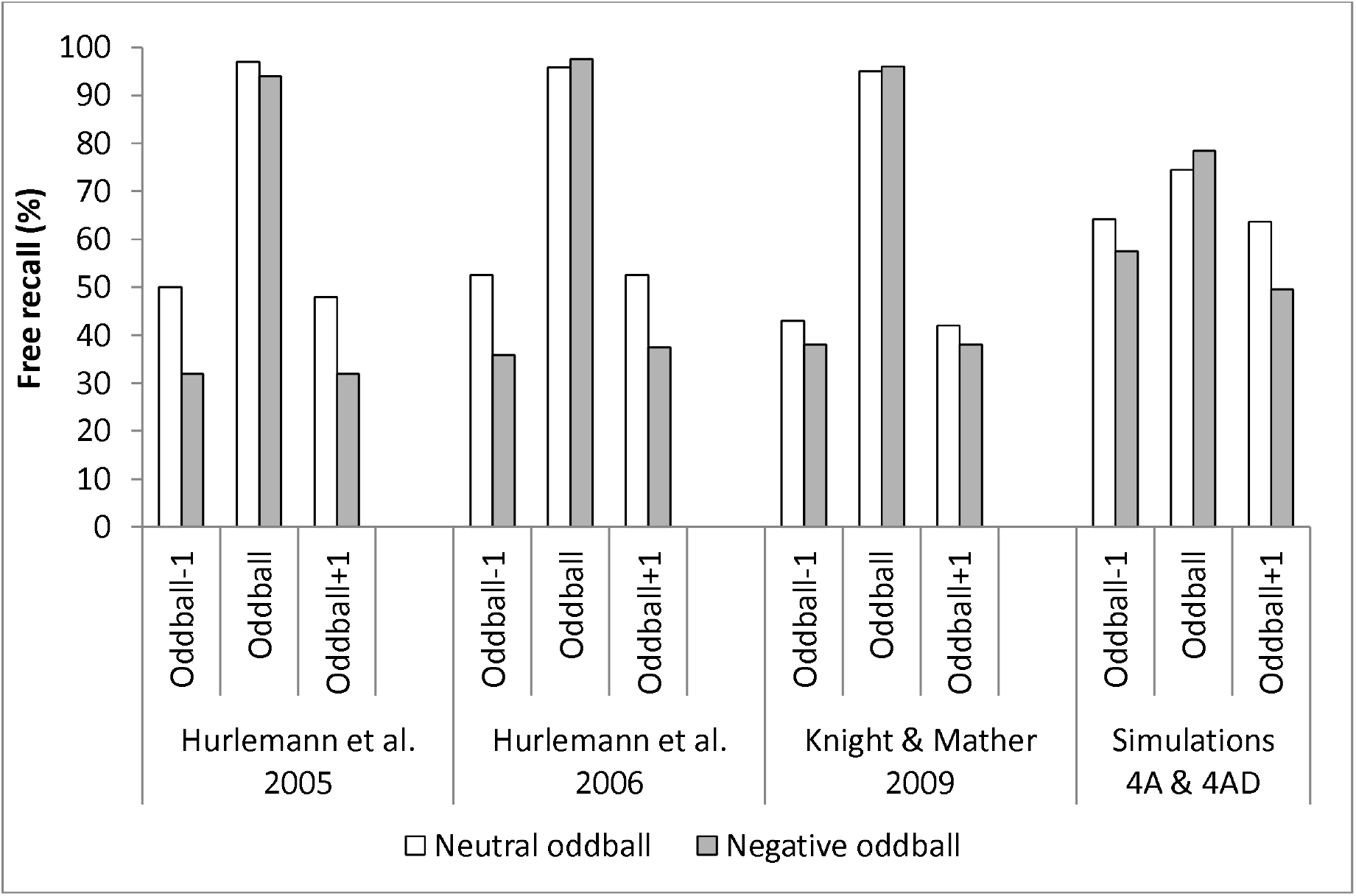
Empirical and simulated results depicting memory for an emotional and neutral oddball and surrounding items. The simulated data refers to the predictions of Simulation 4A (for the neutral oddball) and 4AD (for the emotional oddball).

### 4.2 Comparing emotional and neutral oddball effects: Simulations 4AA and 4AD

Unexpected neutral and emotional items have both been linked to a similar neurobiological mechanism involving temporo-parietal activity and the neurotransmitter noradrenalin (Mather, Clewett, Sakaki, & Harley, 2017; Polich, 2007). To decide which aspect of the emotional oddball effect is specific to emotional oddballs, compared to neutral ones, we refer to a set of empirical investigations of the emotional oddball, unique in that the same procedures were repeated across two laboratories in three different studies (Hurlemann et al., 2005, 2007; Mather & Knight, 2009). Participants in these studies encoded 9 pictures in each list: 8 pictures of everyday objects, and a single emotional or neutral oddball picture in the middle of the list, depicting people in emotional or neutral scenes. A free recall test was administered after a short distractor task. The key findings of these experiments were superior memory for oddballs, both emotional and neutral; and an impairment in memory for the items presented near the emotional oddball, compared to items presented near the neutral oddball (*FIGURE 9*).

We have discussed the ways in which oddballs in eCMR are different from standards. Having a different source, being less related, or more attended, influence memory for the oddball itself; but the disruption to the temporal context occasioned by source context switches is also relevant to memory for nearby positions. As discussed in section 1, CMR assumed that switching between source contexts disrupts the temporal context, making it more difficult to retrieve items that were presented close in time, but have a different source context. This assumption was retained in eCMR, but so far we have assumed that the degree of disruption, controlled by parameter *d*, was not strong. Simulation 4AD considers a more pronounced disruption by increasing *d*. That temporal context disruption plays a larger role in emotional oddball lists compared to mixed lists is consistent with the intuition that a single oddball is more surprising to participants, and therefore results in more unique encoding operations that resemble a task-switch, compared to alternations that occur throughout the entire list, such as when half the list items are emotional and half are neutral. Note that disruptions to the temporal context around an emotional oddball do not impinge on the source context. Thus, the emotional repercussions of viewing a ‘sun-tanning’ model continue to colour the emotional context during the encoding of items that follow the oddball.

The comparison between Simulation 4A and 4AD will help clarify the predictions of eCMR for a neutral and an emotional oddball that are equally attended, but where the emotional oddball (Simulation 4AD) disrupts the temporal context more than the neutral oddball (Simulation 4A). Memory for the ‘emotional’ oddball resembled memory for the ‘neutral’ oddball, but the former caused a decrease in memory for nearby list items (*FIGURE 8*). This pair of simulations therefore captured the empirical findings (*FIGURE 9*), suggesting that the empirical difference between emotional and neutral oddballs (Hurlemann et al., 2005, 2007; Mather & Knight, 2009) has to do with the degree of disruption to the temporal context when the source context switches between emotional and neutral sources.

The temporal organisation of simulated recall suggested that neutral and emotional oddballs were recalled early, in agreement with the predictions the model made for the list composition task, and with empirical data (Elhalal, Davelaar, & Usher, 2014). In the control lists of Simulations 5A and 4AD the item in position 9 was recalled first around 5%-6% of the time. By contrast, the ‘neutral’ oddball in 4A was recalled first 21% of the time, and the ‘emotional’ oddball in 5E was recalled first 53% of the time. Finally, we examined which items the model tends to recall after it recalls the oddball (*FIGURE 8*). This analysis suggests that the disrupted temporal context around the emotional oddball decreased forward transitions to nearby items.

### 4.3 Summary

In contrast to perfectly mixed lists with an equal number of neutral and emotional stimuli, here we consider lists which include a single emotional “oddball” item. It was important to consider the emotional oddball effect, because it appeared potentially challenging to eCMR. Indeed, we saw that with the parameter values used to simulate the list-composition effect we did not obtain an oddball effect. Our aim in this section was to show that eCMR *can* simulate this effect, and offer insights as to the factors that might allow it to do so. We focused on identifying the elements of the oddball effect that were specific to emotional, compared to neutral oddballs. eCMR provided a good qualitative fit to a robust set of results that have been replicated several times in different laboratories.

The simulations conducted here provided novel predictions that can be tested empirically. First, eCMR could capture superior memory for oddballs compared to standards only when oddballs were simulated as attracting more attention than emotional items in mixed lists. To test this prediction future research could manipulate the proportion of emotional items in the list and measure attention empirically by using a divided attention task or EEG recordings (Barnacle et al., 2018; Pottage & Schaefer, 2012; Talmi, Schimmack, et al., 2007). Second, the degree of impairment to recall of items before and after the oddball in lists that included an oddball, and in control lists, was affected by the degree of disruption to the temporal context, not by increased attention to the oddball. The degree to which oddballs disrupt the temporal context can also be measured empirically using subjective time estimations (Block & Reed, 1978; Sahakyan & Smith, 2014).

## 5. Discussion

The work presented in this paper proposes a shift in the way we understand memory for important experiences, those that trigger emotional arousal. It is well-established that some experiences attract attention and are processed preferentially: those with certain features that our species has evolved to prioritise, those previously associated with reward or punishment, and those that appear to promote our current goals. It has previously been thought that the memory traces laid down during the encoding of emotionally-arousing experiences are maintained in a special way so that important events will have the best chance of influencing later fitness-related decisions. Our simulations show that the nature of the memory traces themselves – the increased binding between them and their encoding context, and their inherent associations to other emotionally arousing and semantically related memory traces – may be sufficient to give them an advantage during test, which is amplified in delayed tests, and especially when they compete for retrieval with neutral items. Differences that are already evident at encoding between emotional and neutral items protect and promote them whenever there is a competition for retrieval later on, and give rise to emotion-enhanced memory even without further post-encoding advantages. The dynamics of retrieval determine the magnitude of this advantage and could increase or decrease it. eCMR therefore highlights the exquisite sensitivity of memory to the situation in which an agent finds itself – in eCMR, the retrieval context of the memory test – by providing mechanisms which prioritise the retrieval of experiences that best match that test context across multiple dimensions.

eCMR is the first quantitative model of the emotional enhancement of memory. It explains the improvement in remembering emotionally arousing stimuli as arising from the operation and modulation of retrieved context mechanisms during encoding, maintenance and retrieval. We assumed, in agreement with the consensus in the literature (reviewed, for example, in Mather & Sutherland, 2011; Pourtois et al., 2013) and our own previous findings (Talmi & McGarry, 2012; Talmi, Schimmack, et al., 2007), that stimuli that trigger emotional arousal attract attention obligatorily. Building on previous work, extra attention was described in the model as the strengthening of associations between attended items and their encoding context (Howard & Kahana, 2002a; Lohnas et al., 2015; Polyn et al., 2009), here specifically the source context. eCMR therefore is compatible with the suggestion that emotional stimuli are bound more strongly to their context (Hadley & MacKay, 2006; MacKay et al., 2004; MacKay & Ahmetzanov, 2005), although it specifies that the increased binding is with the sub-region of the encoding context that represented the items’ source. Crucially, eCMR goes beyond previous work, revealing that emotion-enhanced memory is not solely a result of prioritised encoding, but is also the result of retrieval competition, extending to this situation the logic of CMR (Polyn et al., 2009) and its predecessors. The increased association strength between emotional stimuli and their source context, and the shared source context among emotional items, render emotional items more competitive during retrieval even when their semantic cohesiveness is equated with that of neutral stimuli. When the model attempts to retrieve emotional and neutral items that were encoded together in mixed lists, the emotional ones are recalled early and promote each other’s recall, which interferes with and delays the recall of neutral stimuli. When the memory test takes place after a prolonged interval, the model assumes that the temporal context is less diagnostic, so non-temporal context features such as shared emotional context and the oft-stronger semantic associations between emotionally arousing stimuli boost retrieval chances even more. But when emotional items compete against other highly-attended emotional items, the extra attention they receive at encoding does not help very much at retrieval, so that, recall of pure emotional lists is a lot closer to recall of pure neutral lists. In summary, eCMR uses the established mechanisms of retrieved context models to describe emotion-enhanced memory as a consequence the interplay between encoding, maintenance, and retrieval effects.

The data presented here support eCMR as a model of the effect of emotional arousal on immediate free recall. eCMR provided a good qualitative fit for the emotional list composition effect and the emotional oddball effect, phenomena related to differences between the average recall of emotional and neutral stimuli. In the recall of mixed lists the increased average strength of associations between emotional items and their context supported their enhanced recall, while in the recall of pure lists the increase in the strength of average associations between items and their context no longer mattered. In reality, what would matter for recall of pure lists would be the fluctuation in attention that individual participants may allocate to individual items. This realisation now provides an explanation to a finding we reported a few years back, when we used structural equation modelling to relate attention at encoding and free recall performance (Talmi & McGarry, 2012). Participants in that study encoded pure and mixed lists of emotional and neutral pictures under full or divided attention. Performance on the concurrent task worsened in terms of accuracy and reaction time when participants encoded emotional, compared to neutral pictures, regardless of list composition. We used performance on the concurrent task to compute an attention score for every picture (across participants), a measure which correlated with the arousal ratings each picture received. Intriguingly, we found that in free recall of mixed lists arousal ratings predicted recall directly, an effect which was not mediated by attention; but in pure lists the effect of arousal on memory was completely mediated by attention. The current modelling explains these findings because a little more attention to particularly arousing pictures in a pure list can certainly boost retrieval success, but its impact will be drowned out in mixed lists when all emotional items are attended a lot more than all neutral items.

A list with one emotional oddball is also, in a way, a mixed list. eCMR captured the emotional oddball effect too, under the assumption that an oddball attracts more attention than an emotional item in a list with many other emotional items – an assumption that should be investigated empirically. Without that assumption the recall of oddballs was poorer than the recall of control items; but increased attention to the oddball resulted in good memory for it, as well as a decrease in memory for items in nearby positions. Empirical data suggested that only the latter was specific to emotional oddballs, such that anterograte and retrograde amnesia around the oddball only occur because an emotional oddball disrupts the temporal context more than a neutral oddball - a novel prediction. This prediction can be tested empirically through retrospective time estimations, which are sensitive to the number of different patterns that are encoded into episodic memory (Faber & Gennari, 2015) and are thought to index the magnitude of drift of temporal context (Block & Reed, 1978; Sahakyan & Smith, 2014). Recently, evidence emerged (Johnson & Mackay, n.d.) that emotional oddballs disrupt the temporal context. They presented participants with a list of 6 words. At the end of the list participants were asked to estimate the duration of the last word, which was either a taboo word or a neutral word. Participants estimated the duration of the taboo word as longer than the duration of the neutral word. More direct evidence for the predictions of eCMR that emotional oddballs disrupt the temporal context more than neutral oddballs will require comparing the taboo word to a neutrally-valenced oddball word.

eCMR also captured well the dynamics of recall. The stronger association between emotional stimuli and their context in eCMR increases the chances that emotional items will win the retrieval competition early in recall, so the model predicted earlier recall of emotional stimuli studied in mixed lists and the earlier output of oddball stimuli. This prediction has also been confirmed empirically, as shown here for the list composition effect using a number of different datasets, and reported in the literature on the emotional oddball (Elhalal et al., 2014). When there are more than one of them in the list, emotional and semantic associations between emotional stimuli help them promote each other recall, and therefore the model predicted above-chance transitions based on the emotional context of stimuli. This prediction, too, was confirmed, both in previous data where clustering recall around the emotional category was observed (Long et al., 2015; Siddiqui & Unsworth, 2011; Talmi, Luk, et al., 2007), and in new analysis of data from Barnacle et al. (2018). Biasing the model so that recall begins with a more neutral context prevented the model from recalling emotional items first, but they gained the upper hand very quickly. This occurs because when neutral items are recalled they retrieve their context, which helps them promote the recall of items more strongly bound to that context – the emotional items. Emotional items then further promote the recall of items that share their emotional and semantic context. The prediction that emotional items will be recalled better than neutral ones even when the initial retrieval context is biased against them is in keeping with new data we reported here, from an experiment where participants encoded mixed lists, and then were instructed which item type to recall. More emotional items were recalled than neutral ones both in the first recall test *and* in the second recall test, where the instructions switched to the other item type. In summary, this investigation shows a multiply determined effect of emotion on retrieval dynamics, which together with enhanced attention at encoding gives rise to the robust emotional memory enhancement in immediate free recall of mixed lists.

Our simulations showed that eCMR predicts increased retrieval success when the context of encoding and test is similar. This occurred when memory for emotional (neutral) items that were encoded in mixed lists improved when the source context of the test was biased towards emotional (neutral) items. This is reminiscent of state-dependent memory effects – the established improvement of memory performance when participants are in the same ‘state’ at encoding and retrieval (Smith & Vela, 2001). States can physiological, e.g. when induced through psychopharmacological manipulations, or psychological, for example through mood inductions (Blaney, 1986; Bower, 1981; Ucros, 1989). eCMR predicts enhanced recall when the emotionality of the retrieval state matches the emotionality of the encoding context, suggesting that it may be able to capture state-dependent effects. Although here the dynamics of recall meant that their impact dwindled quickly, so the initial state had only a limited impact on average recall, it could have a larger impact in situations where the original context is less accessible.

### 5.1 Relating eCMR to other models

The model we developed here is useful because it can account for many emotional memory phenomena. Some of the effects the model describes, which have been discussed in previous work in terms of the contribution of emotional arousal, are here revealed as similar to non-emotional effects. The emotional list composition effect is, of course, reminiscent of list composition effects that arise from non-emotional manipulations of item typicality (McDaniel & Bugg, 2008). The increased item-to-context associations, which in eCMR are discussed as a consequence of emotional arousal, may also be true for other atypical neutral categories, such as enacted or generated items. Whether eCMR can account for non-emotional list-composition effects depends on whether additional attention is allocated to each one of the unusual stimuli, and on the semantic associations between all of the items. eCMR can perhaps be seen as an extension and quantification of the item-order account, where various item dimensions that receive attention at encoding can balance out memory impairment stemming from reduced attention to others.

The success of eCMR in modelling the emotional list composition effect can be contrasted with the predictions we can extract from the Scale Invariant Memory and Perceptual Learning model (SIMPLE; Brown, Neath, & Chater, 2007). Neath and Brown (Neath & Brown, 2006) set out to explore the list-composition effect in the recall of short and long words. To do so, they proposed that short words are more distinctive than long words because they are easier to comprehend. Distinctiveness was the main mechanism that allowed SIMPLE to capture the findings that memory for a pure list of short words is better than memory for a pure list of long words, and that in mixed lists of short and long words, memory for short and long words is equivalent, and is at the level of memory for a pure list of short words (Hulme, Stuart, Suprenant, Bireta, & Neath, 2004). Why is that relevant for emotional memory? Because like short words, emotional items are also more distinctive than neutral items, due to the unique cognitive and affective processes they trigger during encoding. If we take this analogy seriously (Brown, personal communication) it would seem that SIMPLE predicts the mirror image of the empirical emotional list composition effect: that memory for memory for pure neutral lists will be decreased, compared to all other conditions, just as memory for pure lists of long words is decreased compared to all other conditions.

The emotional oddball effect is also reminiscent of semantic and perceptual oddball effects. In a striking demonstration, the same oddball effects are observed either when a picture of a nude model is the oddball item within a list of pictures of clothed models (the standard items), or vice versa. Therefore, it is reasonable to ask whether models that were developed to explain the generic oddball effect – including the categorization-activation-novelty model, CAN (Elhalal et al., 2014) and SIMPLE (Brown et al., 2007) - can also explain the emotional oddball effect, and whether eCMR can account for non-emotional oddball effects. We did not conduct a model comparison and cannot therefore comment on whether eCMR is better or worse in capturing oddball effects compared to CAN and SIMPLE. The simulation we reported here suggests that the mechanisms that drive both may be similar, and distinguished only in the degree of attention allocated to the oddball, and the degree to which it disrupts the temporal context of encoding. These dimensions can be crossed in an empirical investigation, for example using emotional and neutral oddballs that attract varied amount of attention and disrupt the temporal context to varying degrees. eCMR did make some unique predictions that could be tested empirically to support it against competing models of the oddball effect. First, eCMR predicted increased retrospective time estimation in oddball compared to control lists, which could be modulated by emotionality. Second, while memory impairment for items nearest the oddball is common to many models, eCMR uniquely predicted that this impairment is not a result of increased attention to the oddball. Ultimately, the success of eCMR compared to other models will be revealed by data that test these novel predictions.

### 5.2 Limitations of the modelling approach and plans for future research

The emotional experiences of our lives are rich with personal meaning that defines us as people and connects us to others and to the cultures we live in. They can be positive or negative, happy or calm, threatening or disgusting, and motivate us to approach, avoid, or otherwise behave in multiple bewildering ways. The controversy around defining emotion is well-known (Izard, 2010) and there is no sign yet for an umbrella theory that can bridge the gaps between the different theoretical positions that give rise to the disparate definitions. eCMR simplifies matter by considering only two aspects of emotional experiences–the semantic association between them and their increased intensity (Bradley & Lang, 1994; Feldman Barrett, 2016; Lindquist, Satpute, Wager, Weber, & Barrett, 2015; Russell, 1980). Yet even arousal was only represented in eCMR inasmuch as it drove enhanced attention, an operationalisation most closely related to the ‘relevance detection’ appraisal (Sander, Grandjean, & Scherer, 2005) than to other theoretical treatments.

One aspect of emotional experience that deserves additional consideration is valence, because there is evidence that negatively-valenced stimuli are remembered better than positive ones even when attention is controlled (Kensinger, 2009a) and that valence may modulate some of the effects we discussed, including the relationship between attention and memory in mixed lists (Talmi, Schimmack, et al., 2007) and the effect of oddballs on preceding items (Hurlemann et al., 2005). Future research should test whether the empirical results that we drew on here replicate with positive, arousing stimuli. eCMR can be extended to represent a valence dimension of context, if such results imply that this is necessary. In that case, it would be important for future research to control the semantic associations within and across valence categories, because there is evidence that positive stimuli are judged to be more similar to each other than negative ones (Koch, Alves, Krüger, & Unkelbach, 2016).

The concept of an emotional dimension of similarity, which has been so useful in eCMR, also deserves further research. Our previous work (Talmi & Moscovitch, 2004) inspired researchers to equate the semantic relatedness of the emotional and neutral stimuli they use. The problem with this approach, which we have used extensively ourselves, is that when stimuli are emotional, semantic relatedness ratings may reflect a different construct than they do when stimuli are neutral. Judgements of similarity may be influenced both by semantic associations and by emotional associations related to how stimuli make participants feel. Participants may therefore rate two negative stimuli, for example, two sad facial expressions belonging to two different people, as more related than a sad and a neutral expression belonging to the same person. These open questions notwithstanding, the qualitative mechanisms of binding and competition that we investigate here are robust to quantitative variation in the degree of similarity across items.

eCMR was confined to the simulation of free recall, and in that, like other models, it neglects influences that arise from the nature of the memory test. The advantage of emotional stimuli that we highlighted here would, however, influence any test that benefits from memory for the temporal context of encoding. Although items may not compete with each other in a recognition test as they do in free recall, the decision to classify a memory trace as a ‘Remember’ response is inherently linked to the extent to which the encoding context is retrieved at test. This may explain why emotion enhances measures of recollection more than familiarity (reviewed by Yonelinas & Ritchey, 2015). The integration of emotion into retrieved context models help us decipher an intriguing, recent dataset, where neutral stimuli that were encoded after a block of emotional stimuli were recollected better than neutral stimuli encoded before a block of emotional stimuli, before a block of neutral stimuli, or after a block of neutral stimuli (Tambini et al., 2017). We comment on this study because as in the emotional list composition effect, here, too, the key comparisons are between conditions that include only neutral items and those that mix emotional and neutral ones, and neutral memory changed based on the presence of emotional items. Following principles of retrieved context models, when emotional stimuli retrieve their temporal context during the recognition test, this might help items that share their encoding context retrieve it during the test. This could increase the recollection of neutral items that are studied close after, but not before, the emotional items, because only those share the temporal context of emotional items. In addition, it is likely that participants were still aroused when they encoded neutral items after an emotional block, because the physiological effects of emotional arousal linger for many minutes (Dickerson & Kemeny, 2004), resulting in shared source context as well; and that the arousal state will have increased attention participants allocated to retrieved memory traces (Mather et al., 2015). Therefore, there are a number of reasons compatible with eCMR that could account for the findings, but these speculations could only be examined when eCMR is extended to recognition memory tests.

Another major avenue for future development of eCMR is to simulate the effect of emotion on associative memory. The main challenge is to understand the fact that when participants are re-presented with an emotional item that they have encoded earlier, they are better at retrieving some aspects of the context of encoding, with a consequent increase in accurate Remember judgments, but not other aspects of the context, such as the orienting task that was performed during encoding, or other items that were encoded close in time and space (Bisby & Burgess, 2013; Madan et al., 2012, 2017; Sharot & Yonelinas, 2008; Yonelinas & Ritchey, 2015). These latter results appear to challenge eCMR because of the key role that increased item-context binding plays in the model. Retrieved context models cannot, at present, provide a comprehensive account for associative memory, although there has been preliminary attempts to do so (Davis, Geller, Rizzuto, & Kahana, 2008; Howard, Jing, Rao, Provyn, & Datey, 2009). Because we did not simulate Remember-Know or cued-recall tests, eCMR cannot directly speak to these points. Yet it is important to consider that the predictions of eCMR depend on the interplay of encoding and retrieval effects, and that, crucially, participants encode all aspects of the context of emotionally-arousing items differently than they do the context of neutral items. For example, an emotional element of a complex stimulus is thought to deprioritise other elements of the stimulus (Mather et al., 2017; Mather & Sutherland, 2011). When an emotional item itself captures attention, this takes away from the attention allocated to other stimuli in the same scene (Kensinger, Garoff-Eaton, & Schacter, 2007; Riggs, McQuiggan, Farb, Anderson, & Ryan, 2011). Even the effort participants spend on integrating the emotional item with another member of the same pair is altered (Murray & Kensinger, 2012). It would be important to clarify how these altered encoding processes influence the parameters in eCMR before we can relate our results to findings from the associative memory literature.

Finally, retrieved context models do not have a dedicated consolidation module (but see Sederberg et al., 2011). The core claim is that at the algorithmic level of explanation, time-dependent interference and context changes and reinstatement are sufficient to explain the behavioural phenomena that are typically attributed to consolidation. Retrieved context models are models of the information processing considerations that give rise to memory performance, which therefore can be implemented in various ways neurobiologically. Thus, we by no means intend to deny the contribution of mechanisms such as replay or time- or sleep-dependent change in the brain structures that maintain memory traces and even the very structures of synapses that store individual memories (McKenzie & Eichenbaum, 2011). These are simply outside the scope of our model. Future, more neurobiologically-minded work might clarify how the brain might implement the computations proposed in eCMR, and conversely the eCMR computations might inform the investigation of the neurobiological mechanism. The step we took in section 3, where we ask how delay might change the nature of memory traces and the operation of retrieval mechanisms, has been inspired by behavioural and neurobiological findings about the way that memories are transformed with time delay, but is similarly blind to the actual mechanism that implement such transformations. Instead, the possibility that an increased reliance on non-temporal dimensions of similarity renders memories more schematic with time can be best tested by its fit to behavioural data, even if it is fundamentally inspired by findings that older memory traces rely less on the hippocampus and more on the neocortex (Moscovitch et al., 2016). Crucially, because eCMR can explain both immediate and delayed effects of emotion, it is the first model of emotional memory that encompasses both parsimoniously, but at the cost of a disconnection from the implementation level of explanation.

### 5.3 Concluding remarks

We know that memory performance is not simply a read-out of encoding and maintenance operations, but a reconstruction on the past in the service of current goals (Schacter & Addis, 2007). The current modelling suggests new directions for understanding the adaptive role of emotional memory enhancement in guiding behaviour. These effects have long been understood in terms of prioritizing which individual items to encode and maintain, but situating them within retrieved context models sheds new light on how emotional effects on retrieved memories might support decision making. We have already noted the striking formal correspondence between item-context associations (as inferred from recall tasks and embodied in retrieved context models) and predictive world models that have been separately proposed (and verified; Momennejad et al., n.d.) to underlie the evaluation of candidate actions in decision tasks (Gershman et al., 2012). In brief, the stored learned set of item-context associations amounts to a predictive world model that connects items, situations or events to others they tend to predict. In a decision task, these links can be retrieved at choice time to support a mental simulation of the likely consequences of candidate actions (Shohamy & Daw, 2015). In this framework, the new effect here of emotional modulation of these associations in eCMR would serve to highlight emotionally salient consequences, tending to give them a larger effect on one’s deliberations. This, in turn, formally relates these emotional memory enhancements to a set of computational mechanisms and empirical effects (e.g., Huys et al., 2012) that have been proposed to underlie both adaptive control of prospection and maladaptive dysfunction, such as rumination and worry (Eldar et al., 2016; Huys, Daw, & Dayan, 2015) in disorders of mood.

The core claim of retrieved-context theory, that the conditions of retrieval crucially influence which items are recalled, was extended here to memories that are have a personal, emotional meaning. The important role of the retrieval stage in accessing these memories resonates particularly well with the motivations that drive much emotional memory research: to relieve the suffering that sometimes results from aversive emotional memories, on the one hand, and to extract neutral information about events that are dominated by emotional experiences, on the other. The applied clinical and forensic value of our modelling approach stems from its potential to inspire psychological manipulations to alter the accessibility of emotional and neutral memories, even long after the original experiences.

## Authors note

This work was supported by the Wellcome Trust [105610/Z/14/Z] and the Royal Society [IE160027]. We thank R. Fox, R. El-Nagib, G. Barnacle, E. McManus, M. Slapkova and K. Lutor for data collection and processing. We thank M. Moscovitch, J. Caplan, and K. Norman for their helpful comments. Part of this paper was presented at the Context and Episodic Memory Symposia (2017, 2018) and Learning and Memory (2018). A pre-print of this manuscript was published on BioRxiv.

## APPENDIX 1: DETAILS OF METHODS IN TALMI ET AL.’S (2007) EXPERIMENT 2, SIMULATED IN SECTION 2

In simulations 1-3 we use three variants of eCMR to capture the average recall data from the emotional and related-neutral conditions in Talmi et al., 2007, Experiment 2. The semantic coherence of the stimuli used in that experiment was matched, first, by selecting neutral items that depicted domestic scenes, and second, by equating the semantic similarity ratings of emotional and neutral stimuli obtained in a separate rating study.

In the rating study an independent sample of 13 participants were presented with four sets of 40 pictures. Each set included 40 pictures: 10 emotional negative, 10 related neutral pictures which were all depictions of domestic scenes, 10 random neutral pictures, and 10 positive pictures. The pictures were drawn from the IAPS (Lang & Bradley, 2007) and supplemented by pictures from the Internet obtained using Google image search. Participants were given instructions explaining what semantic relatedness meant, contrasting it to perceptual relatedness, and rated all possible pairs within a subset on a 1 (unrelated) - 7 (highly related). Reliability analysis of ratings within each picture type, across the 4 sets (all negative-to-negative ratings, etc.) resulted in the exclusion of one participant. For the purpose of the 2007 study, several subsets of five pictures from these sets were selected, and a semantic relatedness score was computed for each picture in the subset based on its relatedness to the other 4 pictures in the same subset. Each list in the main experiment included three such subsets. The same pictures were used across the pure and mixed list conditions. The pure lists included 3 subsets of emotional negative pictures; 3 subsets of related neutral pictures; or 3 subsets of randomly selected neutral pictures. The mixed lists included one subset from each of these categories. There was no significant difference between the average semantic relatedness of the emotional and the related-neutral pictures, but the relatedness of random-neutral pictures was lower. The order of pictures in a list was randomised for each participant.

In that experiment two groups of participants studied either 3 pure lists or 3 mixed lists of 15 experimental pictures. Experimental pictures were preceded by two buffer stimuli of the same type (or one emotional and one neutral, in mixed lists), and followed by a minute-long arithmetic distractor task, after which participants recalled the pictures by describing the content of the pictures they have seen in writing. Each recall output was scored as ‘correct’ if two independent raters agreed that the description matches that of one, and only one, of the pictures in the study set. In the rating study participants were presented with subsets of 5 pictures; all pictures in a subset were drawn from the same category (emotional negative, related neutral pictures which were all depictions of domestic scenes, random neutral pictures). The pictures were drawn from the IAPS (Lang & Bradley, 2007) and supplemented by pictures from the Internet obtained using Google image search. Participants rated all possible pairs within a subset on a 1-7 scale for their degree of semantic relatedness. A semantic relatedness score was computed for each picture in the subset based on its relatedness to the other 4 pictures in the same subset. Each list in the main experiment included three such subsets. The same pictures were used across the pure and mixed list conditions. The pure lists included 3 subsets of emotional negative pictures; 3 subsets of related neutral pictures; or 3 subsets of randomly selected neutral pictures. The mixed lists included one subset from each of these categories. There was no significant difference between the average semantic relatedness of the emotional and the related-neutral pictures, but the relatedness of random-neutral pictures was lower. The order of pictures in a list was randomised for each participant.

## APPENDIX 2: DETAILS OF METHODS FOR THE INSTRUCTED RECALL EXPERIMENT, REPORTED IN SECTION 2

We manipulated retrieval of mixed lists of negatively valenced emotional and neutral pictures by asking participants to recall emotional or neutral items first (the ‘emotional first’ and ‘neutral first’ conditions), comparing the results to a ‘natural’ mixed recall where no such instructions were given. Three groups of 18 participants took part in this experiment for course credit. They were 18-35 years old and were screened for neurological and psychiatric history. Each group studied 3 mixed lists of 10 negative and 10 neutral pictures, presented in a random order. The first list was always a practice list, and included unrated practice pictures, used in the practice portion of the Talmi et al. (2007) study. Each of the two experimental lists included the ‘emotional negative’ and the ‘related neutral’ pictures from one of the sets produced in the rating study reported in Talmi et al. (2007; describe above in **APPENDIX 1**). A semantic relatedness score was computed for each picture in the list based on its relatedness to the other pictures in the same set. The semantic relatedness of the negative and related-neutral pictures in each experimental list was equivalent. Note that the semantic relatedness score can be computed in multiple ways; if we only consider the relatedness of the negative and realted-neutral pictures (those that were actually included in the experiment), the relatedness of negative pictures was slightly but significantly higher than the relatedness of related-neutral pictures.

Participants were informed that they will be viewing negative, arousing pictures and neutral pictures of domestic scenes, and that the order of recall would vary. The first recall in all groups was ‘natural’; data from this practice list was discarded. The next two lists were allocated to the same between-participant condition, either natural, emotional first, or neutral first.

When participants encode lists of stimuli that include emotionally arousing ones, the post-encoding emotional context is invariably more emotional than it was before the list was studied. This emotional coloration of the time-of-test context cue before any item is actually retrieved could render emotional stimuli more accessible than a context of a delayed retrieval test, biasing recall to commence with the retrieval of an emotional stimulus first. Therefore, a minute-long arithmetic distractor task separated the study and the picture free recall task to bring the emotional sub-region of the test context back to neutral. The ‘natural’ mixed recall group were given 3 minutes to recall all of the pictures at any order. After those 3 minutes have elapsed, participants were invited to continue recall for another 3 minutes, although most participants could no longer recall anything more at that point, and data for that group was aggregated across these two test periods. The ‘emotional first’ group was given 3 minutes to recall the emotional pictures (first recall test). After 3 minutes, they were asked to recall the neutral pictures for another 3 minutes (second recall test). The ‘neutral-first’ group was asked to recall the neutral pictures first (first recall test). After 3 minutes, they were asked to recall the emotional pictures for 3 minutes (second recall test).

